# Probing protein ubiquitination in live cells

**DOI:** 10.1101/2022.02.13.480252

**Authors:** Weihua Qin, Clemens Steinek, Ksenia Kolobynina, M. Cristina Cardoso, Heinrich Leonhardt

## Abstract

The reversible attachment of ubiquitin governs the interaction, activity and degradation of proteins whereby the type and target of this conjugation determine the biological response. The investigation of this complex and multi-faceted protein ubiquitination mostly relies on painstaking biochemical analyses. Here, we employ recombinant binding domains to probe the ubiquitination of proteins in living cells. We immobilize GFP-fused proteins of interest at a distinct cellular structure and detect their ubiquitination state with red fluorescent ubiquitin binders. With this ubiquitin fluorescent three-hybrid (ubiF3H) assay we identified HP1β as a novel ubiquitination target of UHRF1. The use of linkage specific ubiquitin binding domains enabled the discrimination of K48 and K63 linked protein ubiquitination. To enhance signal-to-noise ratio, we implemented fluorescence complementation (ubiF3Hc) with split YFP. Using in addition a cell cycle marker we could show that HP1β is mostly ubiquitinated by UHRF1 during S phase and deubiquitinated by the protease USP7. With this complementation assay we could also directly detect the ubiquitination of the tumor suppressor p53 and monitor its inhibition by the anti-cancer drug Nutlin-3. Altogether, we demonstrate the utility of the ubiF3H assay to probe the ubiquitination of specific proteins and to screen for ligases, proteases and small molecules controlling this posttranslational modification.

## INTRODUCTION

Protein ubiquitination is a highly conserved posttranslational modification which involves the concerted action of E1, E2 and E3 enzymes to ultimately ligate the carboxyl terminus of ubiquitin to a lysine residue of the selected protein target^1–3^. Proteins can be modified at one or multiple lysine residues with a single ubiquitin or ubiquitin chains formed through one of their lysine residues (K6, K11, K27, K29, K33, K48 and K63) or the N-terminal methionine residue (M1). The K48-linked ubiquitin linkage predominantly signals for proteasomal degradation, whereas the K63-linked linkage is mainly involved in non-degradative processes such as DNA repair and NF- κB signaling^2, 3^. Due to technical challenges and difficulties in the detection of the other types of ubiquitin chains, less information is available about their functions. Since E3 ligases widely control protein levels and function, it becomes increasingly important to identify and characterize their specific targets in order to comprehend their unique contribution to complex regulatory protein networks.

The major approach for substrate identification relies on the physical interaction between a ubiquitin E3 ligase and its substrates, using methods such as yeast two-hybrid (Y2H), protein microarrays and biotin-dependent proximity labelling (BioID)^4–6^. The ubiquitination of substrates has been directly monitored with donor/acceptor fluorophore pairs and sophisticated fluorescence resonance energy transfer (FRET) and fluorescence lifetime imaging (FLIM) techniques^7, 8^. Likewise, ubiquitin conjugation was detected with bimolecular complementation of fluorescence or luminescence^9, 10^ and was used for genetic screens^11^. An alternative approach is to assay for altered protein stability upon chemical or genetic interference with an E3 ubiquitin ligase of interest as ubiquitinated proteins are targeted to proteasomal degradation^12, 13^, which could also be monitored by bioluminescence energy transfer (BRET) in PROTAC treated cells^14^. Still, the gold standard in the identification of ubiquitination targets is mass spectrometry but, as a prominent biological function of this modification is proteasomal degradation, the fleeting abundance of modified target proteins often limits their detection.

To enrich for low abundance peptides from ubiquitinated proteins, antibodies that recognize the ubiquitin remnant motif Lys-e-Gly-Gly (diGly), which is exposed upon tryptic digestion of ubiquitinated proteins, have been developed for global proteomic applications^15–17^. In addition, substrate trapping approaches based on polyubiquitin-binding domain fusions have been generated for the isolation of polyubiquitinated proteins from cell extracts^18, 19^. In contrast to antibodies, these ubiquitin-binding domains, which show different binding affinities for distinct ubiquitin linkages^20, 21^, can also be fused to fluorescent proteins to detect different biological ubiquitin signals^22–24^. Similarly, GFP-tagged ubiquitin has been used to visualize free and linked ubiquitin in cells^25^. While these methods detect general ubiquitination levels and proteome-wide changes^26, 27^, they do not allow to monitor ubiquitination of specific targets in live cells with spatio- temporal resolution.

The detection of specific protein ubiquitination in live cells requires a genetically encoded probe that specifically binds but ideally does not interfere with biological function. Here, we generated recombinant probes consisting of tandem ubiquitin association (2UBA) domains, or ubiquitin interacting motifs (2UIM) or a Npl4 zinc finger (NZF) fused to mCherry. These probes could detect local changes in protein ubiquitination, e.g. at cellular DNA repair foci. To detect the ubiquitination level of specific proteins we immobilized them at defined subcellular sites in a ubiquitin fluorescent-three-hybrid (ubiF3H) assay. The combination of the three ubiquitin probes could reveal the ubiquitination status of specific proteins in live cells. To enhance the signal-to-noise ratio we implemented and validated a protein complementation assay (PCA) that only generates a fluorescent signal if the protein of interest (POI) is ubiquitinated. This versatile tool set is suitable for high-throughput screens to identify E3 ligases and ubiquitin proteases in essentially any organism and to monitor changes over time and throughout the cell cycle.

## RESULTS

### Generation of a genetically encoded ubiquitin probe

Studies of protein ubiquitination mostly rely on biochemical endpoint assays using immunoprecipitation, western blot (WB) analyses and mass spectrometry. To investigate protein ubiquitination in live cells we generated a genetically encoded fluorescent probe based on a naturally occurring ubiquitin binding domain. RAD23 is a DNA repair protein that interacts with the proteasome and shows ubiquitin linkage independent binding for K48- and K63-linked polyubiquitin chains^21, 28, 29^. We fused two ubiquitin-binding domains from RAD23 in tandem with GFP (GFP-2UBA) (Fig. 1A and Extended Data Fig. 1A). To test the binding and precipitation efficiency, we transiently expressed HA-tagged ubiquitin (HA-Ub) together with GFP-2UBA in HEK293T cells and analyzed the co-precipitated ubiquitinated proteins with an HA antibody. Compared with the GFP control, GFP-2UBA was efficient in the specific precipitation of ubiquitinated proteins (Fig. 1B).

**Fig. 1:**
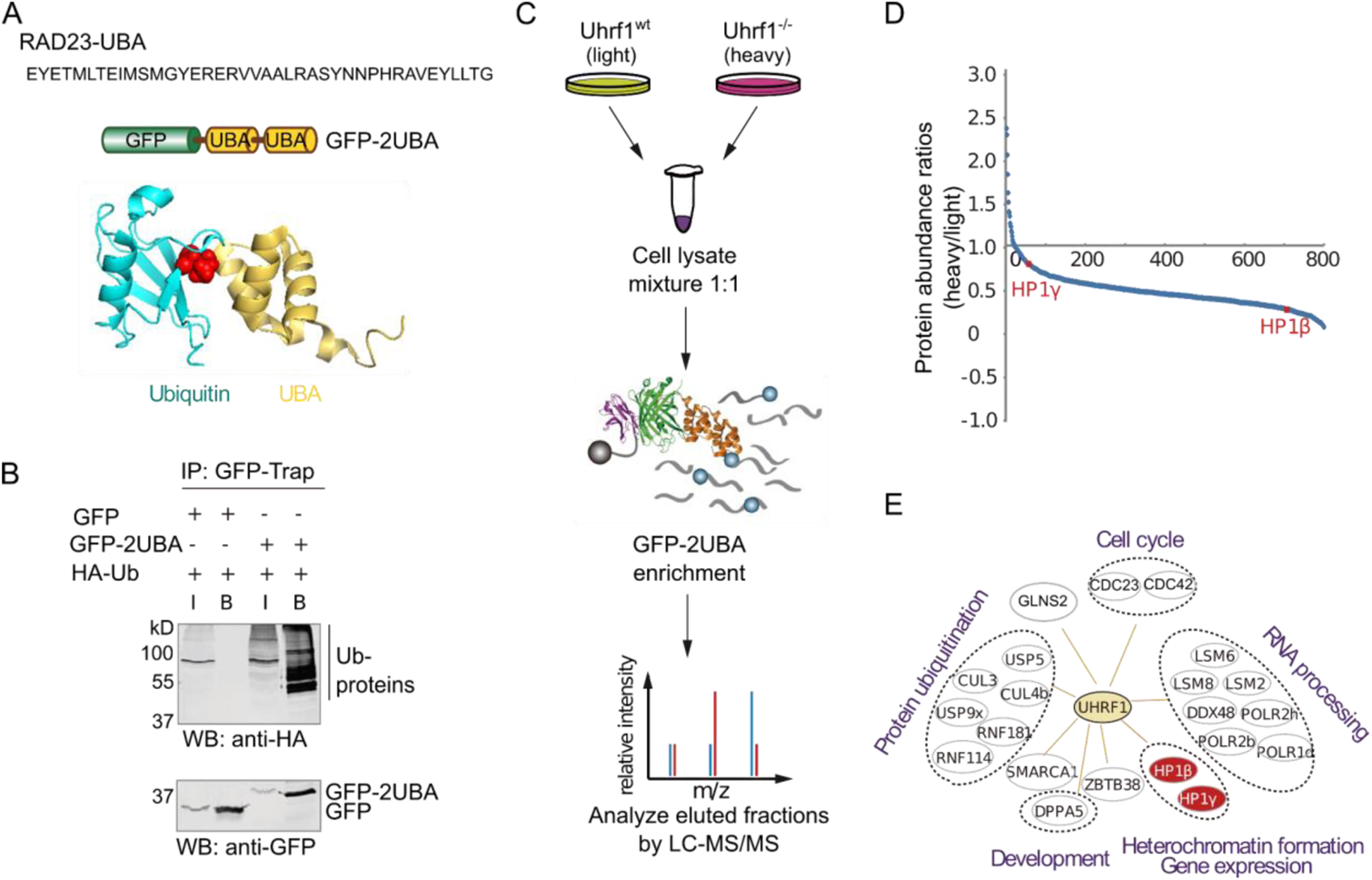
A ubiquitin binding domain co-precipitates with ubiquitin and detects ubiquitination in living cells. (A) Structure and schematic representation of the UBA fluorescent fusion protein as well as the sequence of the UBA domain used to purify ubiquitinated proteins. The structure of the UBA domain (yellow) bound to ubiquitin (cyan) is based on PDB ID code 1WR1. Residue Ile44 at the binding interface of ubiquitin is highlighted in red. (B) Immunoprecipitations with the GFP-Trap from HEK293T cells expressing HA-Ub and GFP-2UBA were probed with an anti-HA antibody to detect coprecipitated ubiquitinated proteins. I and B denote input and bound fractions, respectively. (C) Workflow for identification of UHRF1 ubiquitination targets by SILAC-MS. In this experiment, wt and *Uhrf1*-deficient mESCs were labeled with “light” and “heavy” SILAC medium, respectively. Equal amounts of nuclear extracts from “light” and “heavy” mESCs expressing GFP- 2UBA were mixed for immunoprecipitation. Ubiquitinated proteins were purified with the GFP- Trap and analyzed by LC-MS/MS. See also Extended Data Fig. 1B. (D and E) Heavy/light ratio distribution of all proteins quantified. In total, ∼800 proteins were identified and quantified using the SILAC-based proteomic approach. Target proteins not or less ubiquitinated in *Uhrf1*-deficient cells were selected and grouped according to their gene ontology terms in E.

The RING E3 ubiquitin ligase UHRF1 (also known as Np95 or ICBP90) controls DNA methylation by recruiting the maintenance DNA methyltransferase DNMT1 to hemimethylated DNA substrates^30–33^. To screen for UHRF1 dependent ubiquitination targets, we combined this GFP- 2UBA pull down approach with quantitative mass spectrometry analysis by comparing wild-type (wt) and *Uhrf1*-deficient mouse embryonic stem cells (mESCs) (Fig. 1C). Briefly, wt and *Uhrf1*- deficient mESCs transfected with a GFP-2UBA expression construct were grown in “light” or “heavy” SILAC medium. Equal amounts of nuclear extracts from “light” and “heavy” mESCs were mixed and incubated with the GFP-Trap. Following incubation and washing steps, bound proteins (Extended Data Fig. 1B) were separated and analyzed by liquid chromatography tandem mass spectrometry (LC-MS/MS). Analysis of heavy to light ratios identified proteins that are less ubiquitinated in *Uhrf1*-deficient cells (Fig. 1D), suggesting likely candidates for ubiquitination. In total, ∼800 proteins were identified using the SILAC-based proteomic approach (Supplementary Table S1). UHRF1 target proteins related to development, cell cycle, RNA processing and heterochromatin formation pathways are listed in Fig. 1E and Supplementary Table S2. In our further analysis, we focused on the heterochromatin proteins HP1β and HP1γ as they represent potential new links of UHRF1 and epigenetic regulation.

### Visualizing the ubiquitination of specific proteins in live cells

To monitor the ubiquitination of selected POIs we combined the recombinant ubiquitin probe with our previously developed F3H assay^34^ to develop a ubiquitin fluorescent-three-hybrid (ubiF3H) assay. In this assay, GFP fusion proteins are anchored at a defined subcellular structure like, e.g., the nuclear envelope or a *lac* operator (*lac*O) array inserted in the genome by GFP binding proteins (GBP) and visible as a spot of enriched GFP fluorescence in the nucleus. The ubiquitination of GFP fusion proteins was detected with a mCherry-tagged tandem UBA fusion protein that accumulates at the *lac*O spot if the immobilized GFP fusion proteins are ubiquitinated. The ubiquitin probe can detect ubiquitination changes of POI that are regulated by E3 ligases and deubiquitinases (DUBs) (Fig. 2A). We selected the tandem Ch-2UBA fusion as it showed better binding to the test substrate (GFP-Ub) than the simpler Ch-UBA (Extended Data Fig. 1C and 1D). Based on the mass spectrometry results (Fig. 1D), we focused on the ubiquitination of HP1 proteins. While GFP-HP1β and GFP-HP1γ showed accumulation of Ch-2UBA at the *lac*O spot, no ubiquitination was detected for GFP-HP1α (Fig. 2B). The results are in agreement with immunoprecipitation and WB analyses showing that, in contrast to GFP-HP1γ and RFP-HP1α, GFP-HP1β is strongly ubiquitinated (Extended Data Fig. 2A).

**Fig. 2:**
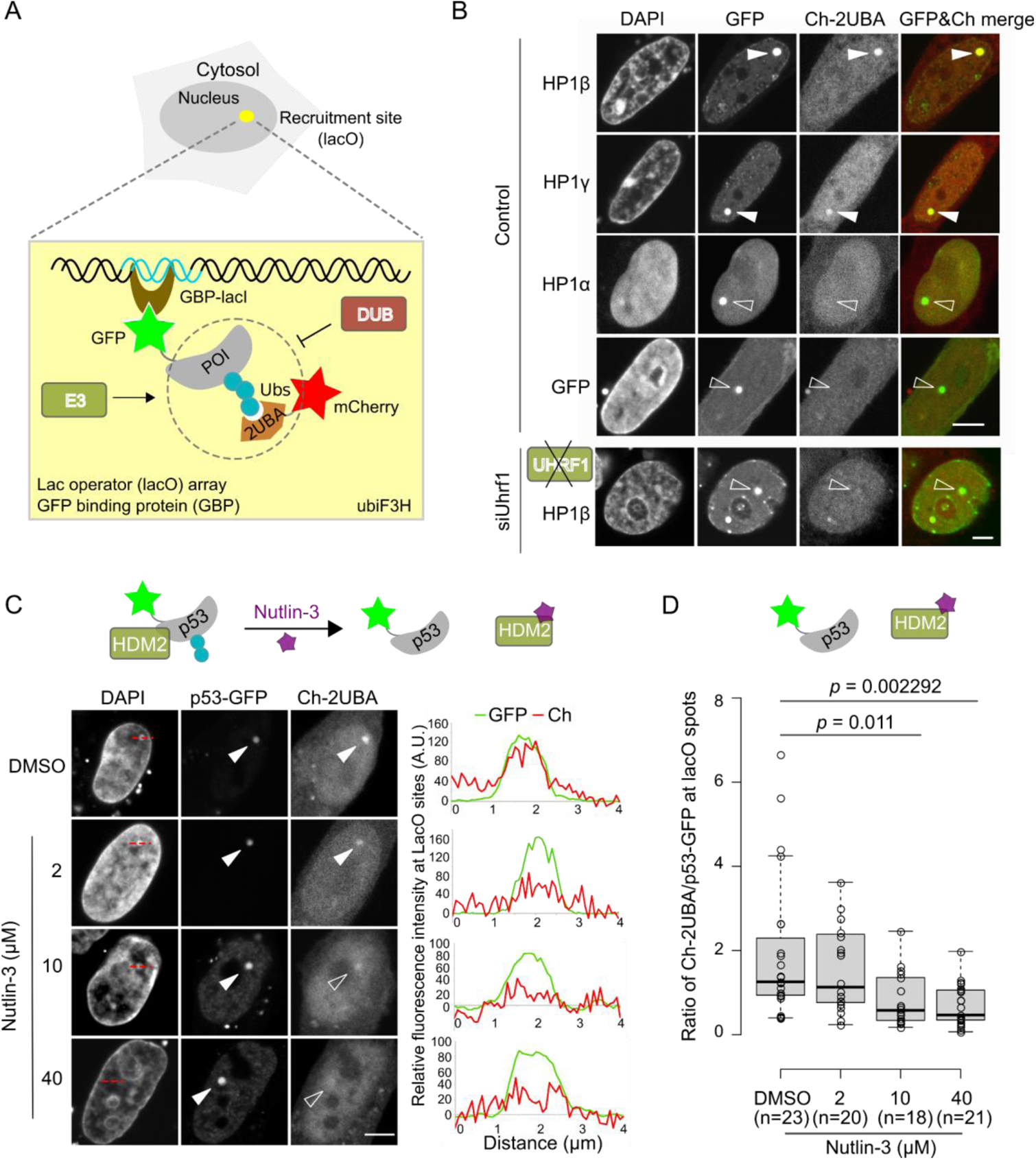
Visualization of the ubiquitination of specific proteins. (A) Schematic representation of the ubiF3H assay to monitor specific protein ubiquitination in live cells. GBP is fused with lacI that accumulates at a *lac* operator (*lacO*) array within the cell. This complex recruits to that particular spot GFP-tagged proteins of interest (GFP-POI). Ubiquitinated GFP-POI can be detected by the Ch-2UBA and visualized using fluorescent microscopy. (B) Ubiquitination of all three HP1 proteins was assayed by ubiF3H assays in BHK wt and compared with *Uhrf1*-knockdown cells (see Extended Data Fig. 4B). The GFP-HP1 fusion proteins were co-expressed with Ch-2UBA in BHK cells. The Ch-2UBA accumulation at *lacO* spots reflects ubiquitination of GFP fusion proteins highlighted with filled arrowheads and not detectable ubiquitination is indicated with open arrowheads. Scale bars: 5 µm. (C) Representative images showing the disruption of HDM2 mediated ubiquitination of p53-GFP by Nutlin-3 at 0 (DMSO, control), 2, 10 and 40 µM. Line scans along lacO spots (red dot lines in the DAPI channel) are shown on the side. Scale bar: 5 µm. Schematic representation of the disruption of p53-GFP and HDM2 interaction by Nutlin-3 is shown above. (D) Quantification of the intensity ratio of Ch-2UBA to p53-GFP at the *lacO* array. Unpaired student t-tests were performed, and *p*-values indicated.

To investigate the binding dynamics of Ch-2UBA in live cells, we performed fluorescence loss in photobleaching (FLIP). We found that the binding of Ch-2UBA to ubiquitinated HP1β is very transient (Extended Data Fig. 3). This transient binding of the ubiquitin probes enables monitoring protein ubiquitination in live cells without interference with biological function.

To identify potential ubiquitination sites, we aligned the three HP1 protein sequences and found three lysine residues at the C-terminus of HP1β not present in HP1γ and HP1α (Extended Data Fig. 2B). Interestingly, these three lysine residues are also part of the recognition sequence KxxxK of the ubiquitin protease USP7 (Extended Data Fig. 8A)^35, 36^. The deletion of the last six amino acids including this KxxxK sequence (GFP-HP1β^delC^) caused a clear reduction in HP1β ubiquitination (Extended Data Fig. 2C).

### Identification of E3 ligases that ubiquitinate proteins of interest

To identify E3 ligases responsible for the ubiquitination of specific proteins we combined our live cell assay with genetic depletion or ectopic expression approaches. We focused on the ubiquitination of HP1β and investigated whether UHRF1 is the E3 ligase, as indicated by the MS screen (Fig. 1E). We first performed *in vitro* ubiquitination assays, in which we immobilized GFP- HP1β on GFP-Trap beads and incubated with recombinant ubiquitin-activating enzyme (E1), ubiquitin-conjugating enzyme (E2), HA-Ub and increasing amounts of His-UHRF1. WB analysis showed clear ubiquitination of GFP-HP1β with increasing amounts of His-UHRF1 (Extended Data Fig. 4A), demonstrating that UHRF1 acts as a ubiquitin E3 ligase for the modification of HP1β. We then performed ubiF3H assays in *Uhrf1*-knockdown cells and found a clear reduction of Ch- 2UBA at the GFP-HP1β spot (Fig. 2B and Extended Data Fig. 4B), similar to DNMT1, H3 and PAF15 (Extended Data Fig. 4C), the known ubiquitin substrates of UHRF1^31, 32, 37–40^.

As an additional example, we chose the tumour suppressor p53, one of the most studied proteins in cancer research. The ubiquitin E3 ligase HDM2 ubiquitinates p53 and thereby targets it for degradation^41^. The cancer drug Nutlin-3 had been developed as a specific inhibitor to disrupt the p53-HDM2 interaction^42^, and thereby prevent p53 ubiquitination (Fig. 2C). With our Ch-2UBA probe, we clearly detected ubiquitination at the p53-GFP spot in control cells (DMSO) (Fig. 2C). The accumulation of Ch-2UBA at *lacO* spots, however, significantly decreased in the presence of more than 10 µM Nutlin-3 indicating reduced ubiquitination of p53 (Fig. 2C and 2D). These results show that our assay is well suited to identify E3 ligases responsible for the ubiquitination of specific proteins and to directly screen for inhibitors in live cells.

### Probes discriminating K48 and K63 ubiquitin linkages

To distinguish the most abundant K48 and K63 ubiquitin linkages (Fig. 3A), we fused ubiquitin interacting motifs (2UIM) from USP25^43^ and the NZF from TAK1 binding protein 2 (TAB2)^24, 44^ with mCherry to generate specific ubiquitin probes (Ch-2UIM and Ch-NZF) (Extended Data Fig. 5A). The 2UIM specifically recognizes K48 ubiquitin linkage via binding the proximal ubiquitin with UIM2 and to the distal one with UIM1^45^, while the NZF binds the Ile44 patch of both proximal and distal ubiquitin to specifically recognize the K63 ubiquitin linkage^44^ (Fig. 3B, Extended Data Fig. 5A).

**Fig. 3:**
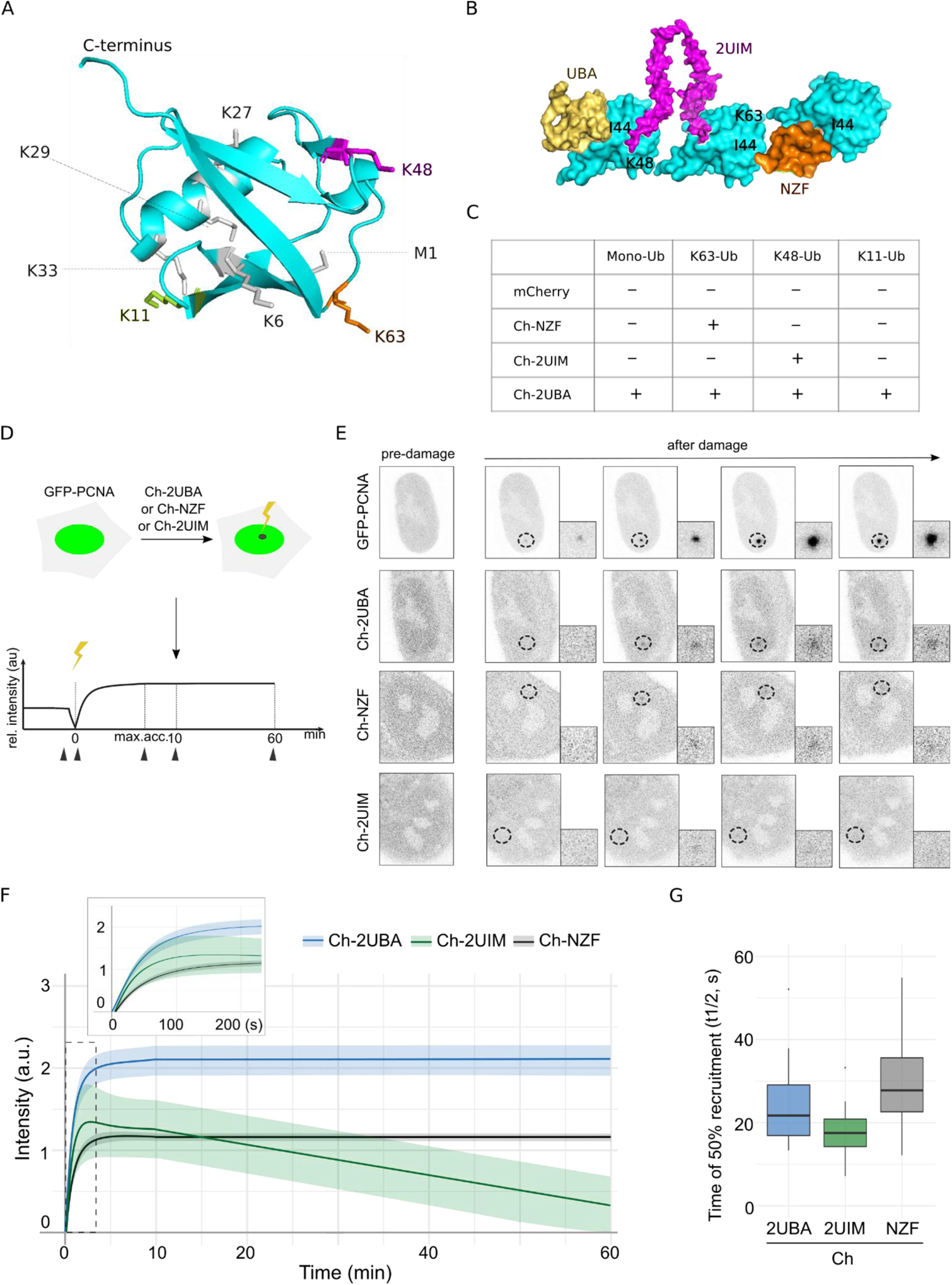
**Discriminating K48 and K63 ubiquitin linkages in live cells. (**A) The structure of ubiquitin (cyan, PDB ID code 1UBQ) and its seven lysine residues for the linkage formation. All seven lysine residues (K6, K11, K27, K29, K33, K48, and K63) and the N- terminal amino group (M1) residing on different sides of the molecule are labeled. (B) Schematic representation of K48 and K63 ubiquitin linkages recognized by UBA, 2UIM and NZF (see details in Extended Data Fig. 5A). (C) Ubiquitination detections of GFP tagged PAF15, DNMT3A, p21, Cyclin B1, p53 and H2A using different ubiquitin probes. Representative images are shown in Extended Data Fig. 5B. The binding preference of ubiquitin probes is summarized according to the quantifications shown in Extended Data Fig. 6. (D) Schematic representation of ubiquitin probes used to monitor protein ubiquitination in live cells at DNA damage sites. (E) Representative images corresponding to the time points indicated in D with black arrowheads. DNA damage was locally induced by microirradiation (black dotted circle) and zoomed images are shown. (F) Ubiquitin probes recruitment curves displayed as mean ± SEM for two or three independent experiments after DNA damage. For each experimental condition at least 25 cells were used. The dotted region is shown as a magnified insert to better illustrate the recruitments of ubiquitin probes within 200 seconds after DNA damage. (G) Times of 50% recruitment of ubiquitin probes at DNA damage sites represented as boxplots. Middle line depicts median value among the cell population.

We used these ubiquitin probes to analyze the ubiquitination of PAF15 (mono-Ub)^38, 39^, DNMT3A (K63-Ub)^46^, p21 (K48- and K63-Ub)^47, 48^, Cyclin B1 (K11-Ub)^49^, p53 (K48-Ub)^50^ and H2A (mono-and K63-Ub)^51, 52^ (Fig. 3C, Extended Data Fig. 5B, 6). The binding preferences detected with Ch- 2UIM and Ch-NZF are summarized in Fig. 3C and are consistent with the mostly biochemistry-based analyses cited above. These results show that the two recombinant probes, Ch-2UIM and Ch-NZF, specifically detect K48- and K63-linked ubiquitin chains, respectively, while Ch-2UBA (Extended Data Fig. 5A) shows a linkage independent binding.

### Detecting protein ubiquitination in live cells with recombinant ubiquitin probes

To test the ability of ubiquitin probes to detect ubiquitination in live cells, we chose DNA damage sites which feature high levels of protein ubiquitination. We locally induced DNA damage in cultured cells stably expressing GFP-tagged proliferating cell nuclear antigen (GFP-PCNA) and ectopically expressing mCherry tagged ubiquitin probes. Repair at DNA damage sites has been shown to involve PCNA, and to elicit ubiquitination and proteasome-mediated degradation^53–55^. To locally induce DNA damage, we irradiated a spot in the nucleus with a focused 405 nm laser. Using live-cell imaging, we monitored GFP-PCNA accumulation as a readout of DNA damage and repair. Accumulation of mCherry tagged ubiquitin probes at the spot of GFP-PCNA accumulation shows that our probes can detect ubiquitination in response to DNA damage (Fig. 3D-3E). By combining these three recombinant probes, we further ascertained the ubiquitination dynamics during DNA repair. We found that Ch-2UIM (K48) showed the fastest association with DNA damage sites (time of 50% recruitment (t1/2): ∼17 sec) versus Ch-2UBA (K48 and K63, t1/2:

∼24 sec) and Ch-NZF (K63, t1/2: ∼30 sec) (Fig. 3F and 3G). In line with previous studies^56–60^, we found that K48-linked polyubiquitination mediates protein degradation and nucleosome eviction at DNA damage sites and facilitates following K63-linked ubiquitination mediated signaling repair pathways. Interestingly, Ch-2UBA and Ch-NZF exhibited stable association with damage sites over one hour, whereas Ch-2UIM dissociated from DNA damage sites after 10 min and its retention time was more variable between cells than of the other two probes (Fig. 3F and 3G). Our observation that the stability of Ch-2UIM binding to the damaged spot was lower indicating its rapid turnover (Fig. 3F and 3G), suggests that the K48 chain formation is short-lived, and less uniform compared to K63^58, 61^ during DNA damage response.

### Enhancing the detection of protein ubiquitination by fluorescence complementation

To further enhance the signal-to-noise ratio and to obtain a single-color readout, we combined this ubiF3H spot assay with a previously described split-YFP^9, 62^ to develop a ubiquitin fluorescence-three-hybrid complementation (ubiF3Hc) assay (Fig. 4A). Briefly, the POI was tagged with the C-terminal half of YFP, whereas 2UBA was tagged with the N-terminal half of YFP. An interaction between POI and 2UBA leading to complementation of both YFP halves yields a functional fluorescent YFP protein, detectable by fluorescence microscopy. YFP shares a high similarity with GFP (Extended Data Fig. 7A) and is also recognized by the GFP binding nanobody GBP^63^. The crystal structure of the GFP-GBP complex shows that the interface extends over both halves of the split YFP and the GBP binds amino acids of the N- and the C-terminal half of the split YFP^64^ to enhance YFP fluorescence (Fig. 4B and Extended Data Fig. 7B, 7C and 7D). As most ubiquitinated proteins are not abundant, the reconstituted YFP proteins are captured at the *lacO* spot using GBPs to improve the signal-to-noise ratio (Fig. 4A). Using the ubiF3Hc assay we confirmed the ubiquitination of HP1β and H3 (Fig. 4C).

**Fig. 4:**
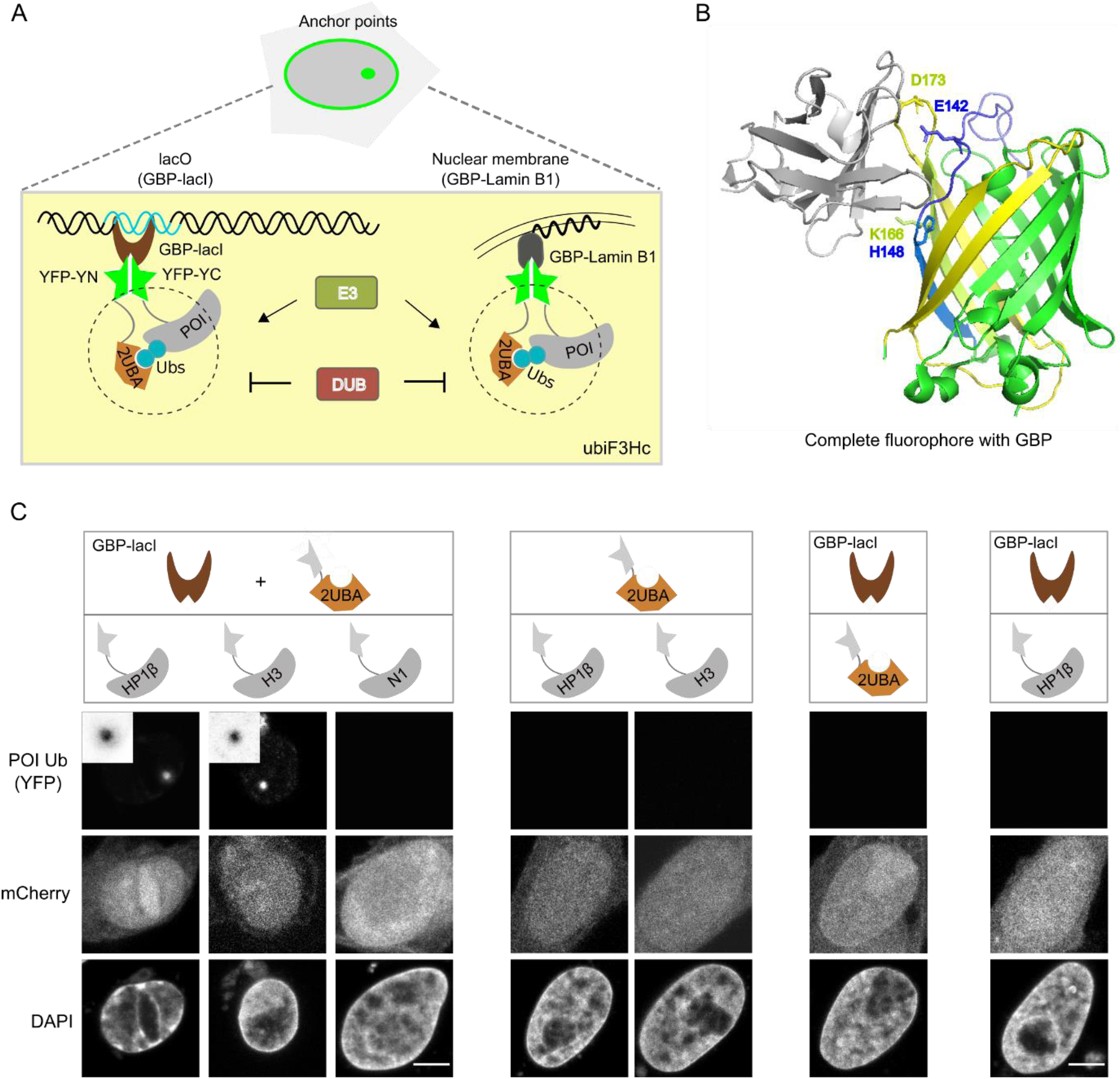
Visualization of the ubiquitination of specific proteins by ubiF3Hc. (A) Schematic representation of a fluorescent protein complementation assay using split YFP fusions, to probe interactions between POIs, fused to the C-terminal half of YFP (YC), and 2UBA, fused to the N-terminal half of YFP (YN). The reconstituted YFP was trapped at the *lacO* array or nuclear membrane by GBP-lacI and GBP-Lamin B1 fusion proteins, respectively. (B) The active reconstituted GFP with GBP protein (PDB ID code 3K1K) is shown and the major amino acids in the binding surface from GFP are highlighted. The GBP protein is labeled in gray. (C) Representative images of H3 and HP1β ubiquitination with the ubiF3Hc assay in BHK cells. The expression of mCherry was used to identify transfected cells. A nanobody fused to the C-terminal half of YFP (N1-YC) and different combinations of those elements were used as negative controls. Scale bar: 5 µm.

We next investigated whether USP7 is the DUB for the deubiquitination of HP1β, as its recognition sequence KxxxK^35, 36^ is found in HP1β (Extended Data Fig. 8A). We observed a significant increase of HP1β ubiquitination in *Usp7*-knockdown cells (Extended Data Fig. 8B, C and D) similar to the positive control H3^65^. Consistently, biochemical analyses showed that GFP-HP1β polyubiquitination was reduced to undetectable levels by co-expression of Ch-USP7, but not the catalytically inactive point mutant Ch-USP7^C224S^ (Extended Data Fig. 8E).

### Monitoring protein ubiquitination along the cell cycle

The ubiF3Hc approach not only improves the signal-to-noise ratio but also frees one color channel for additional readouts to investigate correlations with other cellular processes like, e.g., cell cycle progression by co-expression of RFP-PCNA as S phase marker^66^. While the *lacO* array serves as an efficient anchor point for F3H assays and yields easy to analyze spot signals, it also limits the usage to cells genetically engineered to carry this array. In this regard, besides this *lac*O, also other cellular structures like the nuclear envelope, actin filaments or centrosomes may be used as anchor points for the F3H assays, which allows an easy transfer to other subcellular environments, cell systems and species^34^ (Fig. 4A).

Thus, we applied the ubiF3Hc assay in mESCs using the nuclear envelope as an anchor point to investigate the ubiquitination of PAF15 and H3 during the cell cycle. Recently, we reported that UHRF1 ubiquitinates PAF15 and H3 recruiting DNMT1 for the maintenance of DNA methylation after DNA replication^31^ (Fig. 5A). We compared the ubiquitination timing of PAF15 and H3 during the cell cycle by using the ubiF3Hc assay. We found that PAF15 is predominantly ubiquitinated in the early S phase, while H3 is modified in both early- and middle-S phases (Fig. 5B and 5C). A closer look shows H3 ubiquitination signals throughout the nucleus that seem to accumulate at the rim over time reflecting YFP capture by GBP-Lamin B1 (Extended Data Fig. 9). This result is consistent with our finding that PAF15 is preferentially involved in the methylation of early replicating DNA^39^. We did not detect ubiquitination of PAF15 and H3 in late S phase.

**Fig. 5:**
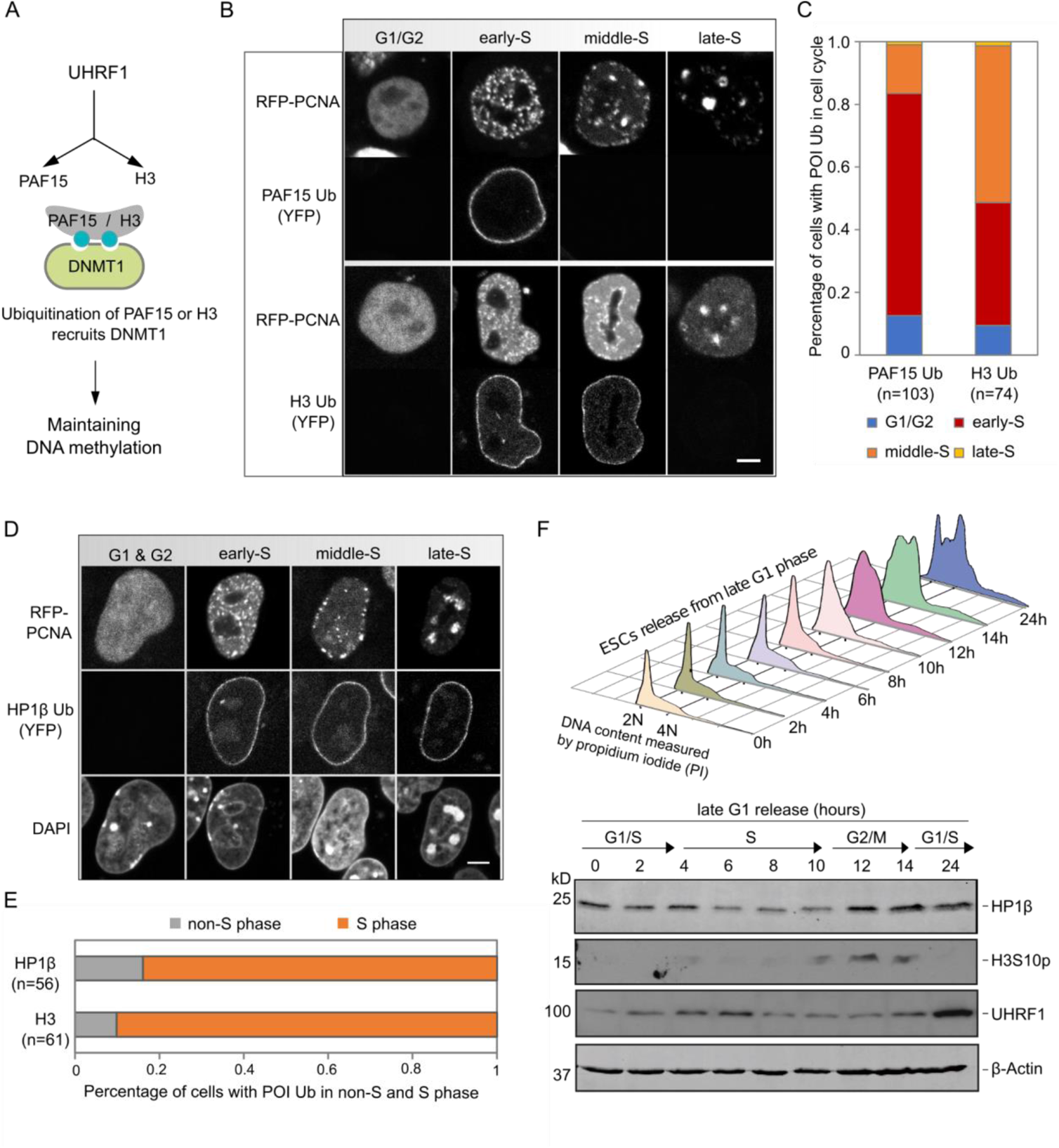
Visualization of cell cycle dependent ubiquitination. (A and B) Illustration of two distinct modes of DNMT1 recruitment after DNA replication for the maintenance of DNA methylation in F. Representative images of cell cycle dependent ubiquitination of PAF15 and H3 in G. Scale bar: 5 µm. (C) Percentage of cells with PAF15 and H3 ubiquitination during the cell cycle. The numbers of cells analyzed (n) are indicated. (D) Representative images of cell cycle dependent ubiquitination of HP1β in mESCs using the nuclear envelope as an anchor point and RFP-PCNA as a cell cycle indicator. Scale bar: 5 µm. (E) Percentage of cells with H3 and HP1β ubiquitination in non-S and S phase. The numbers of cells analyzed (n) are indicated. (F) HP1β and UHRF1 levels in synchronized mESCs released from the late G1 phase were detected by WB. The H3S10p blot was used to highlight the G2 phase, and the actin blot was shown as loading control. Cell cycle profiles of synchronized mESCs are shown above for each time point after release from the late G1 phase.

With the same assay, we found that the ubiquitination of HP1β predominantly occurs in S phase (Fig. 5D and 5E). In parallel, we investigated whether HP1β is subjected to cell cycle dependent regulation by biochemical analyses. In this assay, cells arrested in late G1 phase by mimosine were tested for HP1β abundance at different times after release into the cell cycle. Cell cycle progression was monitored by flow cytometry (Fig. 5F). Consistently, time course analysis of cells released from mimosine G1 arrest showed that HP1β levels decreased in S phase (Fig. 5F), which is consistent with the observed ubiquitination of HP1β (mostly) in S phase. Furthermore, we detected an increase of HP1β level in *Uhrf1*-deficient cells (Extended Data Fig. 10A). These results suggest that UHRF1 ubiquitinates HP1β in S-phase and targets it for degradation.

Thus, our results show that the ubiF3Hc assay can complement and extend biochemical approaches and provide insights into cell cycle dependent changes in protein ubiquitination.

To test the application of ubiF3Hc in monitoring the ubiquitination of specific proteins in live cells, we stably expressed the HP1β fused with the C-terminal half of YFP and transiently expressed the other components. The ubiquitinated HP1β (YFP-HP1β Ub) was recruited to the nuclear envelope (Fig. 6A and Extended Data Fig. 10B). With live cell microscopy, we monitored the ubiquitination of HP1β during cell cycle and consistently observed its ubiquitination in S-phase (Fig. 6B and 6C).

**Fig. 6:**
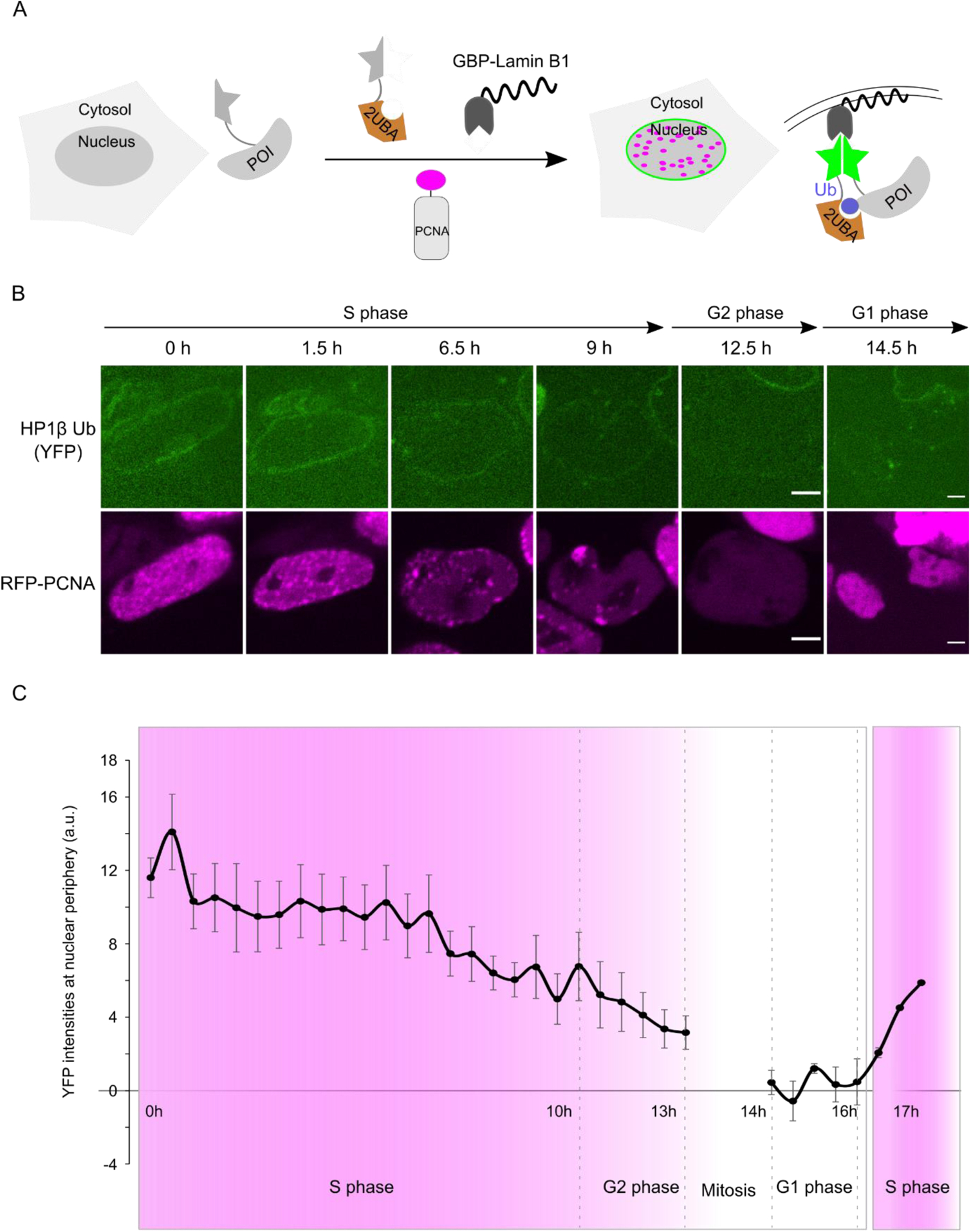
Monitoring the ubiquitination of HP1β in live cells during cell cycle. (A) Schematic outline of the protein ubiquitination monitoring strategy. POI fused with the C-terminal half of YFP was stably expressed in mESCs and other components were transiently expressed to recruit the ubiquitinated protein to nuclear envelope. (B) Live cell series of mESCs stably expressing YC-HP1β and transiently co-expressing components shown in A. Scale bars: 5 µm. (C) Time lapse quantification of the YFP (HP1β-Ub) intensities at the nuclear lamina. Five individual cells were aligned and the mean and the SEM of the YFP intensities were shown. The images of cells at mitosis states are excluded from the quantification. The last two time points were derived from one single cell.

### Identification of targets and inhibitors of ubiquitination in live cells

As the ubiF3Hc assay shows a high signal-to-noise ratio, we next sought to examine whether it is applicable for high-throughput screening in drug discovery. We tested this assay in 96-well plates to automatically visualize and analyze HP1β ubiquitination. In line with the result shown in Fig. 2B and 4C, we observed clear reconstituted YFP signals at *lacO* spots in two independent replicates showing HP1β ubiquitination and demonstrating the robustness of the ubiF3Hc assay (Extended Data Fig. 10C). We further recruited ubiquitinated proteins to major satellites (MaSat) that allows the application of ubiF3Hc in any mouse cell types.

To test the potential of this system for identification of E3 ligases, we stably expressed the YC- HP1β and YC-PAF15, the ubiquitination targets of UHRF1, in mESCs (Extended Data Fig. 10B). We then analyzed their ubiquitination level after knock down of *Uhrf1*. As a control, we analyzed the ubiquitination of PAF15 in UHRF1 ubiquitination defective cells (UHRF1 H730A)^67^. In line with the result shown in Extended Data Fig. 4C, we detected significant reductions of PAF15 ubiquitination in both UHRF1 H730A and *Uhrf1* knocked down cells (Extended Data Fig. 10D). Likewise, we observed a reduction of HP1β ubiquitination in *Uhrf1* knocked down cells (Extended Data Fig. 10D) that is in line with our MS result (Fig. 1D).

To investigate the potential of this ubiF3Hc assay for the identification of specific inhibitors, we chose the well-known tumor suppressor p53 which is ubiquitinated by HDM2 and thus marked for degradation^41^. To restore and boost p53 tumor suppressor activity in cancer therapy, small molecules like Nutlin-3 have been developed to inhibit the ubiquitination by HDM2^42^. We expressed the components of the ubiF3Hc assay for detection of p53 ubiquitination and clearly observed reconstituted YFP signals at *lacO* spots in consistency with Fig. 2C and 2D. We then incubated the cells with 10 µM Nutlin-3 to test the potential of this assay for the identification of specific inhibitors. The reconstituted YFP signals at *lacO* spots significantly decreased indicating reduced ubiquitination of p53 (Fig. 7A and 7B). These results show that our ubiF3Hc assay is suited to detect the ubiquitination of the tumor suppressor p53 and to identify specific inhibitors like Nutlin-3.

**Fig. 7:**
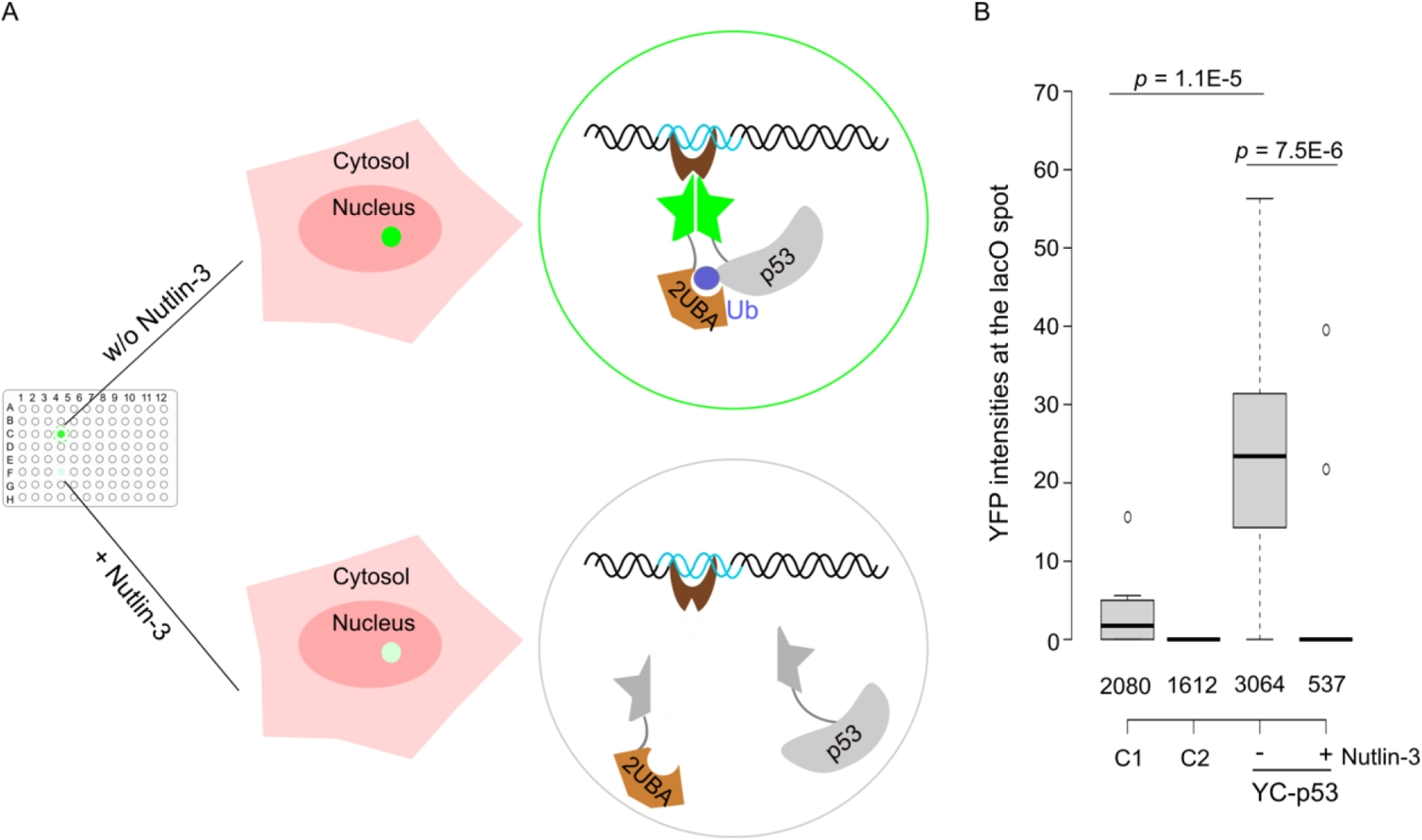
Identification of targets and inhibitors of ubiquitination in live cells. (A) Schematic outline of the detection of p53 ubiquitination in 96-well plates in the presence or absence of Nutlin-3. (B) Box plot representations of intensity ratios of reconstituted YFP and mCherry at *lacO* spots and nucleus, respectively, for p53 ubiquitination in the absence and presence of 10 µM Nutlin-3 (right). C1 (control 1, with N1-YC) and C2 (control 2, without YC-p53). Center lines show the medians; box limits indicate the 25th and 75th percentiles as determined by R software; whiskers extend 1.5 times the interquartile range from the 25th and 75th percentiles; outliers are represented by dots. The numbers of cells analyzed are indicated. Data sets were tested for significance with an unpaired t-test and *p*-values are indicated. Thus, the ubiF3H assay is a robust tool for studying protein ubiquitination in live cells. We could use ubiF3H assays to monitor the ubiquitination of specific proteins in live cells, to identify E3 ubiquitin ligases and proteases, to map the ubiquitination sites and to discriminate different ubiquitin linkages. Most importantly, the ubiF3Hc assay is suitable for studying temporal protein ubiquitination in live cells and high-throughput screens with small molecule or siRNA libraries.

## DISCUSSION

Protein regulatory networks are largely governed by posttranslational modifications and proteasome-mediated degradation, whereby ubiquitination plays a central role. As protein ubiquitination occurs in different configurations and functions, the key to a comprehensive understanding of this dynamic and multifaceted posttranslational modification is the identification of targets with their respective ligases and proteases along with the timing and linkage of this modification. Targets are typically identified by immunoprecipitation with antibodies against the diGly motive remaining after digestion of ubiquitinated proteins as we previously used to identify UHRF1 targets^38^. Here, we co-expressed a GFP-2UBA fusion and enriched ubiquitinated proteins with our nanobody against GFP (GBP) for quantitative MS analysis. We screened UHRF1 dependent ubiquitination targets by comparing *Uhrf1*-deficient versus wt mESCs and identified the heterochromatin protein HP1β as a novel substrate which had not been picked up in our prior study using an antibody against diGly^38^. In general, however, the enrichment with the co-expressed GFP-2UBA was less efficient and low abundant ubiquitinated proteins, like PAF15^38^ were missed. While there are some reports of HP1 ubiquitination^68^ the modification of HP1β by UHRF1 had been missed so far.

The expression of GFP-2UBA allows the detection of ubiquitinated proteins in live cells in general but cannot distinguish which proteins are modified. To detect ubiquitination of specific proteins we developed the ubiF3H assay and expressed GFP fusions with POIs and immobilized them at distinct subcellular structures with the GBP. We used cell lines with a genomic *lacO* array as anchor points to obtain easy to discern spot signals for automated image analysis. To be more flexible and independent from these cell lines, also endogenous structures, like the nuclear lamina, major satellite DNA repeats, actin filaments or centrosomes, may be used as anchor points as previously demonstrated for protein interaction assays^34^. Clearly, the GBP-mediated capture of GFP fusion proteins at a distinct spot also improves the signal-to-noise ratio and the intracellular dynamics seems to be sufficient so that even interactions between mitochondrial proteins could be monitored at nuclear *lacO* spots with the original F2H assay^69^. Also, for convenience we mostly relied on transient transfections but for studies at physiological expression levels, endogenous genes may be tagged as described^70^. This simple ubiF3H assay was not designed for absolute quantification but provides a rapid display of the ubiquitination status of POIs and allows to monitor relative changes in the ubiquitination signal in response to defined manipulations. Thus, we could identify UHRF1 dependent ubiquitination of HP1β and identify the protease USP7 as potential antagonist of this PTM (Figs. 2B, Extended Data Fig. 8 and 10). The UBA domain detects as far as we know all ubiquityl residues and is therefore ideally suited for primary screens with the ubiF3H assay. But as most information of this PTM is encoded in the linkage^2^, i.e. which lysine residues are used for conjugation, we employed more specialized domains for further discrimination. To expand the scope of the ubiF3H assay, we chose the naturally occurring domains 2UIM and NZF that specifically bind K48 and K63 linked ubiquitin chains^43–45^. The comparative use of these three ubiquitin binding domains in the ubiF3H assay makes it possible to discern the most prominent ubiquitin chains in live cells. The ubiF3H assay could be further expanded to identify additional types of ubiquitin chains by using natural or artificial binders like specific affimers that were developed for recognition of K6 and K11/K33 ubiquitin linkages^71^.

The application of fluorescent ubiquitin binders *per se* is limited to monitoring changes in ubiquitination at distinct cellular structures or local processes. FLIP experiments (Extended Data Fig. 3) suggest that the binding of ubiquitin probes is transient and will not interfere with biological functions *in vivo*. Thus, we detected an increase in ubiquitin at focal sites of DNA damage (Fig. 3E) reflecting the known role of ubiquitination in e.g. DNA double-strand repair^72^. The comparison of three ubiquitin chain specific probes showed a faster recruitment of 2UIM at DNA damage sites (Fig. 3F and 3G). This early K48-ubiquitination event at the DNA damage spots is supported by previous publications highlighting, for example, RNF8-mediated K48-linked Ku80 removal^58^, RNF8-mediated K48-linked VCP/p97 and 53BP1 recruitment^59, 60^, and L3MBTL1 degradation^56^. All of these processes happen relatively early after the damaging event and contribute to the formation and stability of repair foci. Some studies have reported the rapid turnover of the K48- linked polyubiquitin chains^58, 61^, which has hindered the study of the K48-linked polyubiquitin chains’ contribution to DNA damage signaling. On the other hand, the long retention times of Ch- 2UBA and Ch-NZF can be explained by K63 polyubiquitin chains’ importance at later DNA damage signaling steps, including the repair pathway choice and subsequent repair^57^.

To enhance the signal-to-noise ratio of the ubiF3H assay, we implemented a fluorescence complementation (FC) approach between POI and ubiquitin reader using split YFP. While this FC approach allows detection of ubiquitination in the natural subcellular environment, it may also spread the signal throughout the cell, depending on the distribution of the POI. Therefore, we co-expressed GBP-Lamin B1 to locally capture the complemented YFP fusions at the nuclear envelope. The binding of GBP is expected to enhance the fluorescence of YFP as was reported for GFP^64^ and to stabilize the complemented YFP as the binding interface spans both halves (Fig. 4B). The local entrapment of the YFP fusions may not only improve the signal-to-noise ratio through the concentration of the signal at a distinct structure but may also slow down degradation. These features of the ubiF3Hc assay enhance the signal but complicate the quantification. Also, as FC depends on spatial proximity and steric fit, the ubiF3Hc assay may, for some POIs, require optimization of linker length and fusion site for the complementing YFP halves. In comparison with the ubiF3H assay, which essentially measures the colocalization of green and red fluorescence, the ubiF3Hc assay requires only a single-color channel (YFP) leaving another channel free for additional readouts like, e.g., the cell cycle phase, as demonstrated with RFP-PCNA (Fig. 5B and 5D).

Previous bimolecular complementation assays relied on the covalent ligation of labeled ubiquitin^9, 10^. Here, we use ubiquitin binding domains, which, by the transient nature of their binding and discriminating specificity, is expected to be more dynamic, less disruptive and more informative. The comparative use of UBA, 2UIM and NZF ubiquitin binding domains in the colocalization based ubiF3H assay to discriminate between monoubiquitination versus K48 and K63 linked ubiquitin chains (Fig. 3) can easily be applied to the complementation based ubiF3Hc assay. Both assays can be further expanded as more specific binding domains are either identified or artificially generated, like. e.g., the aforementioned affimers^71^ making the ubiF3H assays a dynamic and powerful platform for the study of protein ubiquitination in live cells.

One important feature of the ubiF3Hc assay is the improved signal-to-noise ratio achieved by local signal enrichment at distinct subcellular structures like the nuclear envelope or the *lacO* array spot. This worked well for relatively mobile proteins like HP1β with a half recovery time (t1/2) of 2.5 s^73^ but, surprisingly, also for rather immobile proteins like core histone H3 with recovery times in the range of hours^74^. It will be interesting to investigate whether the captured H3 fusion protein stems from a small mobile fraction mobilized by transcription and DNA replication or by ubiquitination itself.

The single color based ubiF3Hc assay is ideally suited to correlate the ubiquitination of specific proteins with other cellular processes like, e.g., cell cycle progression. With RFP-PCNA as a cell cycle marker we found that PAF15 and H3 are preferentially ubiquitinated in S phase (Fig. 5B), which is consistent with the known function of the UHRF1 E3 ligase. A potential limitation of fluorescence based bimolecular fluorescence complementation is its reported irreversibility *in vitro* and *in vivo*^75, 76^. However, we monitored dynamic changes of HP1β ubiquitination during the cell cycle (Fig. 6). This dynamics could be caused by reversible complementation or degradation as described before^77–79^.

UHRF1 is an essential factor for the maintenance of DNA methylation after DNA replication in S phase^30, 33^. On the one hand, UHRF1 controls DNMT1 abundance by polyubiquitination^37, 40^. On the other hand, it mono-ubiquitinates PAF15 and histone H3, and thereby indirectly recruits DNMT1 to replication foci for the maintenance of DNA methylation^31, 32, 39^. Genetic data indicate that PAF15 ubiquitination is preferentially involved in the methylation of early replicating DNA^39^ which fits well with the observed ubiquitination mostly in early S phase (Fig. 5B and 5C). Surprisingly, no ubiquitination was detected for PAF15 and H3 in late S phase, which might point towards a third, still unknown recruiting mechanism or simply reflect limitations in the accessibility in densely packed, late replicating heterochromatin. In this context it is interesting to note that DNMT1 has distinctively slower FRAP recovery kinetics in late S phase^80^. The observation that also HP1β is ubiquitinated by UHRF1 in S phase is novel and gives rise to speculations about possible functions in DNA replication and/or epigenetic regulation which are interesting starting points for future research.

Dysregulation of protein ubiquitination and degradation plays important roles in the development of human diseases^81, 82^. As several E3 ligases have been implicated in human cancer^83^, the specific targeting of these enzymes is presently investigated as a less toxic alternative to current proteasomal inhibitors. However, high-throughput screening for inhibitors and modulators of E3 ubiquitin ligases mostly rely on cumbersome and costly biochemical assays and cell extracts based Ub-detection systems^84–87^. Clearly, the most prominent ubiquitination target in cancer biology is the tumor suppressor p53. Ubiquitination by the E3 ligase HDM2 marks p53 for degradation and keeps its cellular levels low during normal cell cycle progression. Only upon DNA damage p53 levels increase to affect either cell cycle arrest or trigger apoptosis^88^. To boost the activity of mutant p53 in tumor cells small molecules were screened to prevent p53 ubiquitination.

Sophisticated and costly high-throughput screens eventually yielded Nutlin-3, which prevents p53 ubiquitination^42^. We could detect the ubiquitination of p53 with our robust ubiF3Hc assay and monitor a drop in ubiquitination upon addition of Nutlin-3 with a simple optical readout in a multiwell format (Fig. 7). As the ubiF3H assays are cell-based they provide besides the ubiquitination status of POIs also data on cell permeability, bioavailability and toxicity of candidate drugs and should, thus, be well suited for high-throughput and high-content drug screens.

In summary, we present a versatile ubiF3H assay to investigate the ubiquitination of specific proteins in live cells. We demonstrate that this simple assay is well suited to identify new targets, map ubiquitination sites, discriminate different types of ubiquitination, identify E3 ligases, proteases (DUBs) and inhibitors controlling the ubiquitination of specific proteins and monitor changes over time.

## METHODS

### Expression constructs

The coding sequence of the UBA domain of RAD23 (amino acids 158 to 212), the NZF domain of TAB2 (amino acids 663 to 693) and the 2UIM domain of USP25 (amino acids 97 to 140) was amplified using cDNA from mouse E14 ESCs. To generate the GFP-2UBA and Ch-2UBA constructs, a duplicate UBA coding sequence was subcloned into both the pCAG-GFP-IB and the pCAG-Ch-IB vector. The generation of expression constructs for Ch-USP7 (wt, full length), Ch- USP7^C224S^, GFP-H3, GFP-PAF15, GFP-DNMT1, GFP-DNMT3A and HA-Ub was described previously^32, 40, 89^. The human UHRF1 cDNA was ligated into pcDNA3.1 vector with EcoRI and HindII. To generate the GFP-HP1α, GFP-HP1β and GFP-HP1γ constructs, the HP1α, HP1β, HP1β^delC^ and HP1γ coding sequences were amplified using cDNA from mESCs and subcloned into pCAG-GFP-IB vectors.

The DNA sequence coding for full length of p21 and Cyclin B1 was amplified from mouse cDNA by PCR using Phusion high-fidelity DNA Polymerase (New England BioLabs) and cloned in frame into Ch plasmid with AsiSI and NotI restriction endonucleases (New England BioLabs). The mammalian expression constructs for RFP-PCNA, human p53, pGBP-lacI and pGBP-Lamin B1 were described previously^34, 63, 90^.

The duplicate UBA coding sequence (2UBA) was subcloned to fuse with YN (1-154) by a 4× GGSG linker. Protein coding sequences including H3, PAF15, p53 and HP1β carrying AsiSI and NotI restriction cutting sites were subcloned to fuse with YC (155-235) by a 4× GGSG linker.

All constructs (Supplementary Table S3) were verified by DNA sequencing.

### Cell culture, transfection, inhibitor treatment and cell line generation

HEK293T, BHK, HeLa and mESCs were cultured and transfected as described previously^40, 91^, with the exception that Lipofectamine 3000 (Invitrogen) was used for transfection of mESCs. For the *in vivo* ubiquitination assay, transfected HEK293T cells were incubated with medium supplemented with 2 mM N-ethylmaleimide (NEM) for 30 min before harvesting. For Nutlin-3 treatment, before transfection Nutlin-3 was added into the medium and incubated for ∼16 hours. For cell cycle analysis, mESCs were cultured in a medium containing 0.8 mM mimosine^92, 93^ for 24 h. Synchronized cells were then released into the cell cycle by adding fresh medium after washing once with medium. At different time points, samples were harvested for both WB analysis and flow cytometry (Aria II, Becton Dickinson). In brief, cells were washed twice with phosphate buffered saline (PBS), fixed with 70% ethanol for 15 min on ice and finally stained for 40 min at 37°C in solution containing 50 µg/ml propidium iodide (PI), 0.1 mg/ml RNase A, 0.05% Triton X-100. After washing once with PBS, the cell cycle profile was analyzed by flow cytometry.

The human cervical carcinoma HeLa Kyoto cells (ATCC No. CCL-2), HeLa Kyoto GFP-PCNA cells were grown in DMEM medium supplemented with 10% FCS, L-glutamine, and antibiotics at 37 °C in a humidified atmosphere of 5% CO2. HeLa Kyoto cell lines expressing fluorescent PCNA variants were generated^94^ using the Flp-In recombinant system. HeLa Kyoto Ch-NZF, Ch-2UBA, and Ch-2UIM cells were obtained by transfection with the plasmids bearing mCherry gene and NZF, 2UBA, 2UIM genes respectively. Positively transfected cells were selected visually. Cells were seeded on the µ-Dish 35 mm (cat.no 81158, ibidi) in concentration 200.000 cells per dish. Cells were incubated for 24 h after transfection in the humidified atmosphere as described above. Cells stably expressing POI fused with one half of YFP were generated as previously described^95^.

### Identification of UHRF1 targets

SILAC labeling of mouse ES cells was performed at 37°C in ESCs medium supplemented with 100 µg/ml of light (L) or heavy (H) arginine and lysine isotopes, for L: Arg0 and Lys0 (L-arginine and L-lysine, Sigma-Aldrich), for H: Arg10 and Lys8 (arginine-13C6, 15N4 and lysine-13C6, 15N2, Silantes). In addition to the specific lysine and arginine, the completed ESC medium contained knockout DMEM (Silantes), 10% dialyzed serum, 6% knockout serum replacement, 2 mM L-glutamine, 0.1 mM non-essential amino acids, 50 µM 2-mercaptoethanol, 1000 units/ml leukemia inhibitory factor LIF, 1 µM MEK inhibitor PD and 3 µM GSK-3 inhibitor CHIR (2i, Axon Medchem). To assess the SILAC labeling efficiency, cells were cultured in SILAC medium for 10 passages and tested by MS. For identification of the targets, mESCs were first transfected with an expression vector for GFP-2UBA, the immunoprecipitation assay was then conducted as described below with minor modifications. Wt and *Uhrf1*-deficient cells were lysed in buffer containing 150 mM NaCl, 10 mM Tris-HCl (pH7.5), 2.5 mM MgCl2, 2 mM phenylmethylsulphonyl fluoride and 0.5% NP-40, 1x Protease Inhibitor (Serva) and 1 µg/ul DNAase on ice for 30 min and cleared by centrifugation at 4°C. Protein concentrations of cleared cell lysates were measured using the PierceTM 660 nm protein assay kit. Equal amounts of cell extracts were combined and incubated with the GFP-Trap for 2 h at 4°C under gentle rotation. The samples were separated by SDS-PAGE and prepared for LC-MS/MS as described^96^. As a control, cell extracts from wt mESCs (light) expressing GFP were equally mixed with clear cell lysates from *Uhrf1*-deficient mESCs (heavy) expressing GFP-2UBA for immunoprecipitation with the GFP-Trap.

### Live-cell DNA damage assay

Live-cell DNA damage assay was carried out as previously described^97^. Imaging and microirradiation experiments were performed using a Leica TCS SP5II confocal laser scanning microscope (Leica Microsystems, Wetzlar, Germany) equipped with an oil immersion Plan-Apochromat x100/1.44 NA objective lens (pixel size in XY set to 76 nm) and laser lines at 405, 488, 561 and 633 nm. All imaging was conducted in a closed live-cell microscopy chamber (ACU, Olympus) at 37°C with 5% CO2 and 60% humidity, mounted on the Leica TCS SP5II microscope. The emission of GFP and mCherry was captured using the detection range 495-549 and 610- 680, respectively. For standard microirradiation, a preselected spot in non-S phase cells (1 μm diameter) within the nucleus was microirradiated for 0.6 seconds with the laser lines 405, 488, 561 nm laser set to 100%, or for 1.5 seconds with the laser lines 488 and 561 nm laser set to 100%. Before and after microirradiation, confocal image series of one mid nucleus z-section were recorded in 15 seconds intervals.

Photobleaching of mCh-NZF, mCh-2UBA, or mCh-2UIM at previously microirradiated spots was performed using a circular region of interest (1 μm diameter) for 1 s with a 561 nm laser set to 100%. Before and after microirradiation and photobleaching, a confocal image series of one mid-nuclear z-section was recorded in 15 s intervals.

All analysis steps for the confocal microscopy images from microirradiation experiments were performed using ImageJ^98, 99^. Images were first corrected for cell movement and subsequently mean intensity of the irradiated region was divided by the mean intensity of the whole nucleus (both corrected for background) using ImageJ software. For each experimental condition at least 25 cells were used. Half-times for ubiquitin probes accumulation were calculated from the time of microirradiation till maximal accumulation with one phase association (single exponential function: Y = Y0 +( Plateau – Y0) × (1 – e(-K × X).

FRAP data were normalized by pre bleach fluorescence intensity. All fits were performed on averaged normalized FRAP curves and the resulting fit parameters are reported as the mean ± SEM for two or three independent experiments. Curve fitting was done to double the exponential equation.

### Fluorescence loss in photobleaching (FLIP)

Fluorescence loss in photobleaching (FLIP) experiments were conducted using a Nikon TiE microscope equipped with a Yokogawa CSU-W1 spinning-disk confocal unit (50 μm pinhole size), an Andor Borealis illumination unit, Andor ALC600 laser beam combiner (405 nm/488 nm/561 nm/640 nm), Andor IXON 888 Ultra EMCCD camera, Andor FRAPPA photobleaching module, and a Nikon 100×/1.45 NA oil immersion objective. The microscope was controlled by software from Nikon (NIS Elements, ver. 5.02.00). Cells were transfected for 24 hours, plated on bottom 2-well imaging slides (Ibidi) and maintained at 37 °C with 5% CO2 using an environmental chamber (Oko Labs). Pre-bleach images were acquired with the 488 nm and 561 nm laser using 500 ms exposure time with a final pixel size of 130 nm. For each cell, an area of 66 x 151 pixels covering half of the nucleus was bleached with a 561 nm laser which was moved with a dwell time of of 50 μs over the bleaching area and images were acquired every 1.51 seconds. For analysis, the intensity of the spot and the nucleus were manually measured in Fiji for each timepoint and the background was subtracted from the obtained values. Visible spot intensities were normalized by subtracting the respective nucleus intensity and dividing by pre-bleach intensity of the spot. For mCherry controls, the measured mCherry intensities at the visible GFP spots were divided by pre-bleach intensity of the spot. Images of cells with visible drift were discarded.

### ubiF3H and ubiF3Hc assays

The ubiF3H and ubiF3Hc assays were performed as described previously^34^. In brief, mESC or BHK cells containing multiple lac operator repeats were transiently transfected on coverslips and fixed with 3.7% formaldehyde 18 h after transfection. For DNA counterstaining, coverslips were incubated in a solution of DAPI (400 ng/ml) in PBST and mounted in Vectashield. Cell images were collected using a Leica TCS SP5 confocal microscope equipped with Plan Apo 63x/1.4 NA oil immersion objective and lasers with excitation lines 405, 488, 594 and 633 nm, or Nikon TiE microscope mentioned above. To quantify the ubiquitination within the *lacO* spot in the ubiF3H assay, the following intensity was measured by ImageJ and ratios were calculated for each cell: (mCherrySpot − mCherry_Nucleus_)/(GFPSpot − GFP_Nucleus_) in order to account for different expression levels. To quantify the ubiquitination within the *lacO* spot in the ubiF3Hc assay, the following intensity ratio was calculated for each cell: (YFPSpot − YFPNucleus)/mCherryNucleus in order to account for different expression levels.

### High-throughput microscopy and image analysis

The high-throughput microscopy is as previously published with minor differences^34^. Simply, 10^4^ BHK cells plated onto 96-well plates (Greiner Bio-One) were transfected with the plasmids indicated. Before transfection, the medium was supplemented with the Nutlin-3 or DMSO (control). About 16 hours, cells were then fixed with 3.7% formaldehyde and DNA counterstained with DAPI. After cell fixation, images were acquired automatically with an Operetta high-content imaging system at the wide-field mode using a 40x air objective (PerkinElmer). DAPI, YFP and mCherry fluorescent fusion proteins were excited, and the emissions were recorded with standard filter sets.

The images were then analyzed with the Harmony analysis software (PerkinElmer). Briefly, the images were first segmented by intensity and area size according to the DAPI fluorescence using the top-hat method to define the cell nucleus area. The cell population with mCherry signals was considered as transfected cells and chosen for analysis. Then the *lacO* foci were recognized by their intensity in the GFP channel within the nuclear area of cells with mCherry signals. In these cells, the mean intensities of the GFP channel at the *lacO* were recorded, and the ratio of GFP mean intensity at the *lacO* spot to the mean intensity mCherry of the whole nucleus were calculated in order to account for different expression levels.

The images from Extended Data Fig. 10D were first segmented by intensity and area size according to the DAPI fluorescence using the top-hat method to define the cell nucleus area. The cell population with mCherry signals was considered as transfected cells and chosen for analysis. Then the MaSat foci were recognized by their intensity in the 488 nm channel within the nuclear area of cells with mCherry signals. In these cells, the mean intensities of the 488 nm channel at the MaSat sites and nucleus were recorded, and the YFP intensities at MaSat sites were calculated with Eq. (YFPUb = YFPMaSat – YFPnucleus).

### Co-immunoprecipitation (Co-IP)

GFP and RFP-fusion pulldowns using the GFP- and RFP-Trap (ChromoTek) were performed as described^63^. For detection of ubiquitinated proteins by immunoprecipitation, cells were lysed in buffer containing 150 mM KCl, 50 mM Tris-HCl (pH 7.4), 5 mM MgCl2, 1% Triton X-100, 5% Glycerol, 2 mM phenylmethylsulphonyl fluoride and 2 mM 2-mercaptoethanol and 5 mM NEM. After brief sonication, cell lysates were cleared by centrifugation at 4°C for 10 min. Supernatants were incubated with the GFP-Trap beads for 2 h at 4°C under gentle rotation. The beads were then washed three times with lysis buffer and resuspended in SDS-PAGE sample buffer. The anti-HA mouse monoclonal antibody 12CA5 was used for detection of ubiquitinated proteins.

### In vitro ubiquitination assay

The *in vitro* ubiquitination assay was performed as previously described with minor modifications^100^. His- tagged human UHRF1 was purified using Ni-NTA sepharose resin (Qiagen). Recombinant E1 (His-UBE1), E2 (GST-UbcH5b) and HA-Ub were purchased (Boston Biochem). GFP-HP1β from transfected HEK293T cells was immunoprecipitated with the GFP-Trap and incubated at 37°C for 60 min with the complete ubiquitin reaction system consisting of reaction buffer (25 mM Tris-HCl pH 7.6, 5 mM MgCl2, 100 mM NaCl, 1 µM DTT, 2 mM ATP), 200 ng of E1, 200 ng of E2, 500 ng of UHRF1 and 3 µg of HA-Ub. After washing with a wash buffer (20 mM Tris-HCl pH 7.6, 150 mM NaCl and 0.5 mM EDTA), ubiquitination of HP1β was detected using an anti-Ubiquitin antibody (Santa Cruz Biotechnology).

### Western blots

Following separation on SDS–PAGE, samples were transferred onto a nitrocellulose membrane and incubated with indicated antibodies see Supplementary Table S3. Blots were developed with the Pierce ECL western blotting substrate (Thermo Scientific) and scanned by the Amersham™ Imager 600 system.

### siRNA knockdown, RNA Isolation and Quantitative RT-PCR

100,000 BHK cells were plated into a 6-well plate and transfected with 5 nM of siRNA pool (siTOOL) against *Uhrf1* or *Usp7* by Lipofectamine 3000 (Invitrogen). After 24 or 48 hours, cells were harvested for RNA isolation or or kept for further assays. Total RNA was isolated from wt and *Uhrf1*- or *Usp7*-knockdown BHK cells using the nucleospin triprep kit from Macherey-Nagel. 500 ng of total RNA was reverse transcribed with a high-capacity cDNA reverse transcription kit (Applied Biosystems) according to the manufacturer’s instructions. Real-time PCR was conducted using LightCycler® 480 SYBR Green I Master (Roche) on a LightCycler® 480 System (Roche). PCR efficiency and primer pair specificity were examined using a standard curve of serially diluted cDNA and melting curve, respectively. After normalizing to the transcript level of *Usp7* or *Uhrf1*, data was analyzed based on the 2^-ΔΔCt^ method.

## ACKNOWLEDGMENTS

We are grateful to M. Muto and H. Koseki (RIKEN Center for Integrative Medical Sciences, Yokohama) for providing wt and *Uhrf1*-deficient mESCs. We are grateful to Stefan Jentsch (Max Planck Institute of Biochemistry, Munich) for providing the HA-ubiquitin, Jack A. Bates for N1-YC construct and Axel Imhof (Ludwig Maximilians University, Munich) for mass spectrometric analysis. We thank Hartmann Harz (Ludwig Maximilians University, Munich) and the Center for Advanced Light Microscopy (CALM) for support in microscopy. CS is a fellow of the International Max Planck Research School for Molecular Life Sciences (IMPRS-LS). This work was funded by the Deutsche Forschungsgemeinschaft (DFG, German Research Foundation) grants SFB 1361 Project-ID 393547839 to M. C. C. and SFB 1064 Project-ID 213249687 to H.L. and the Bayerische Forschungsstiftung (AZ-1286-17) to H.L..

## AUTHOR CONTRIBUTIONS

W.Q and H.L designed the study and wrote the manuscript. M.C.C. discussed the project and contributed to the manuscript writing. W.Q, C.S. and K.K. performed experiments and prepared Figures. All authors reviewed the manuscript.

## Extended Data

**Extended Data Fig. 1:**
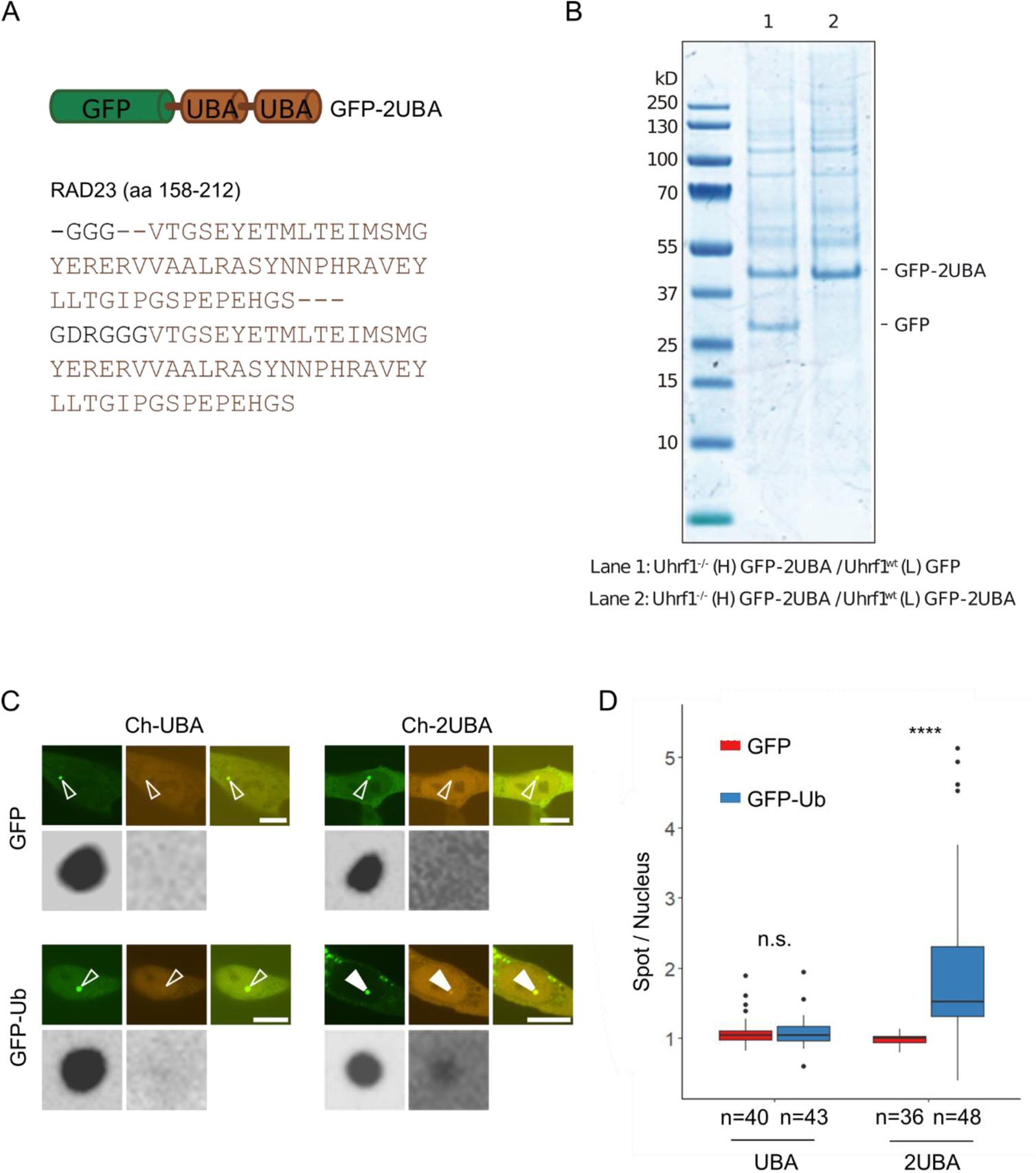
Related to Fig. 1C and Fig. 2A. GFP-2UBA binds ubiquitinated proteins. (A) Two UBA from RAD23 amino acids 158-212 linked with a GDRGGG linker is fused to GFP. (B) GFP-2UBA immunoprecipitations from SILAC labeled mESCs were visualized by Coomassie Brilliant Blue staining. Heavy labeled *Uhrf1*-deficient and light labeled wt mESCs were transfected with a construct coding for GFP-2UBA (Lane 2). Equal amounts of cell extracts from both samples were mixed for immunoprecipitation with the GFP-Trap. As a control, cell extracts from wt mESCs labeled with light amino acids expressing GFP were equally mixed with cell lysates from heavy labeled *Uhrf1*-deficient mESCs expressing GFP-2UBA (Lane 1). Immunoprecipitation was performed with the GFP-Trap, bound fractions were separated by SDS-PAGE, and gels were sliced and subjected to LC-MS/MS analysis. (C and D) Comparison of Ch-UBA and Ch-2UBA binding. The Ch-2UBA accumulation at lacO spots reflects the binding to GFP-Ub highlighted with filled arrowheads and not detectable binding is indicated with open arrowheads. For quantification, Ch signals at the spot were measured and divided by the average signal in the nucleus. Scale bars: 10 µm.

**Extended Data Fig. 2:**
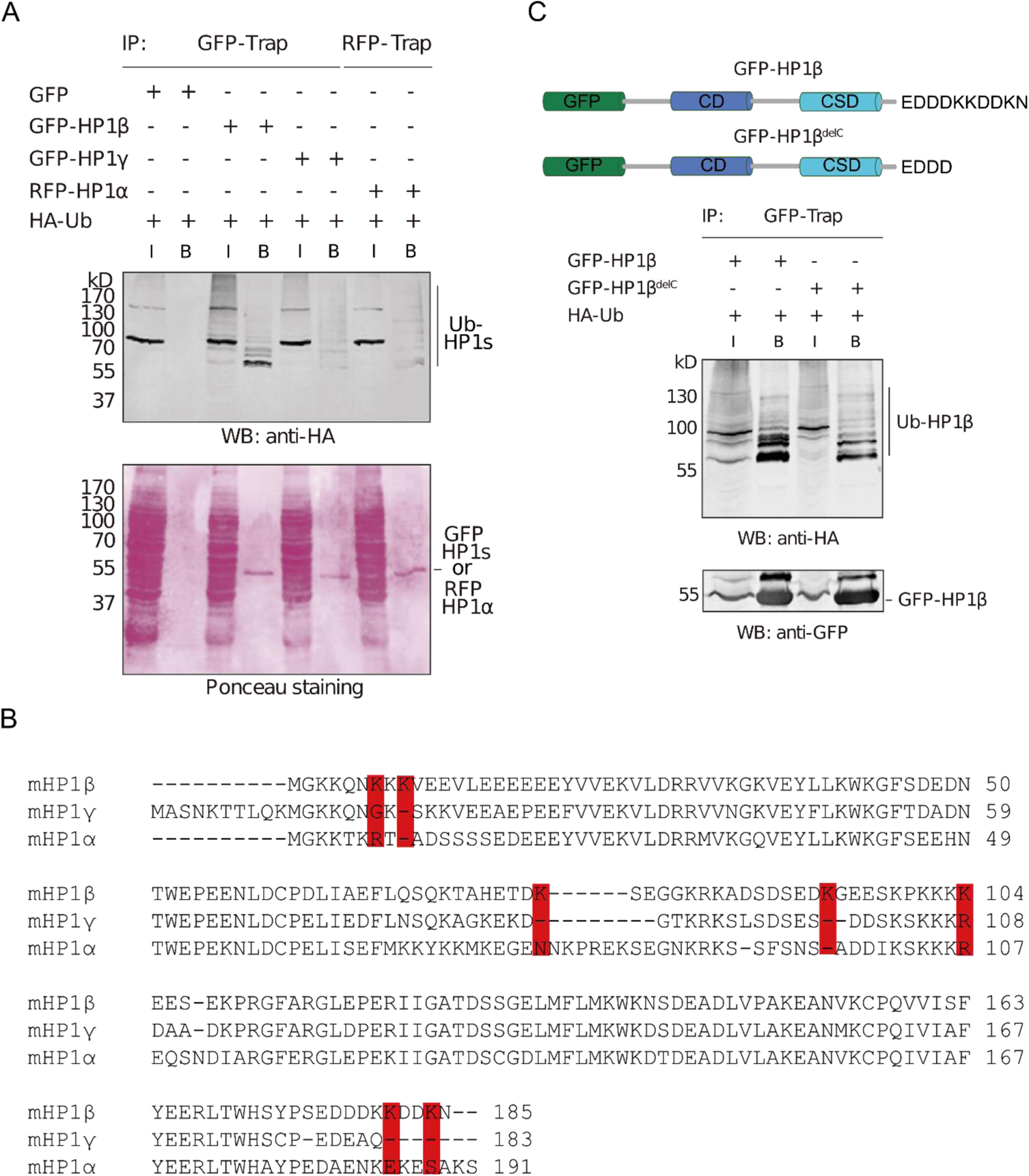
Related to Fig. 2B. Ubiquitination of HP1 proteins and mapping of HP1β ubiquitination sites. (A) GFP- or RFP-Trap immunoprecipitations from HEK293T cells expressing the indicated combinations of HA-Ub, GFP- and RFP-HP1 were probed with an anti-HA antibody. The Ponceau staining was used as a loading control. (B) Alignment of mouse heterochromatin proteins HP1β, HP1γ and HP1α. Lysine residues specific for HP1β are highlighted in red. Accession numbers: HP1β NP_031648.1; HP1γ AAI10377; HP1α AAH04707. (C) Schematic structure of HP1β and its delC mutant and characterization of HP1β deletion mutant to narrow down ubiquitination sites by *in vivo* ubiquitination assay. GFP-HP1β and its mutant were co-expressed with HA-ubiquitin in HEK293T cells and immunoprecipitated using the GFP-Trap. Immunoprecipitations were analyzed with an anti-HA antibody. The delC deletion had a clear effect on the ubiquitination of HP1β.

**Extended Data Fig. 3:**
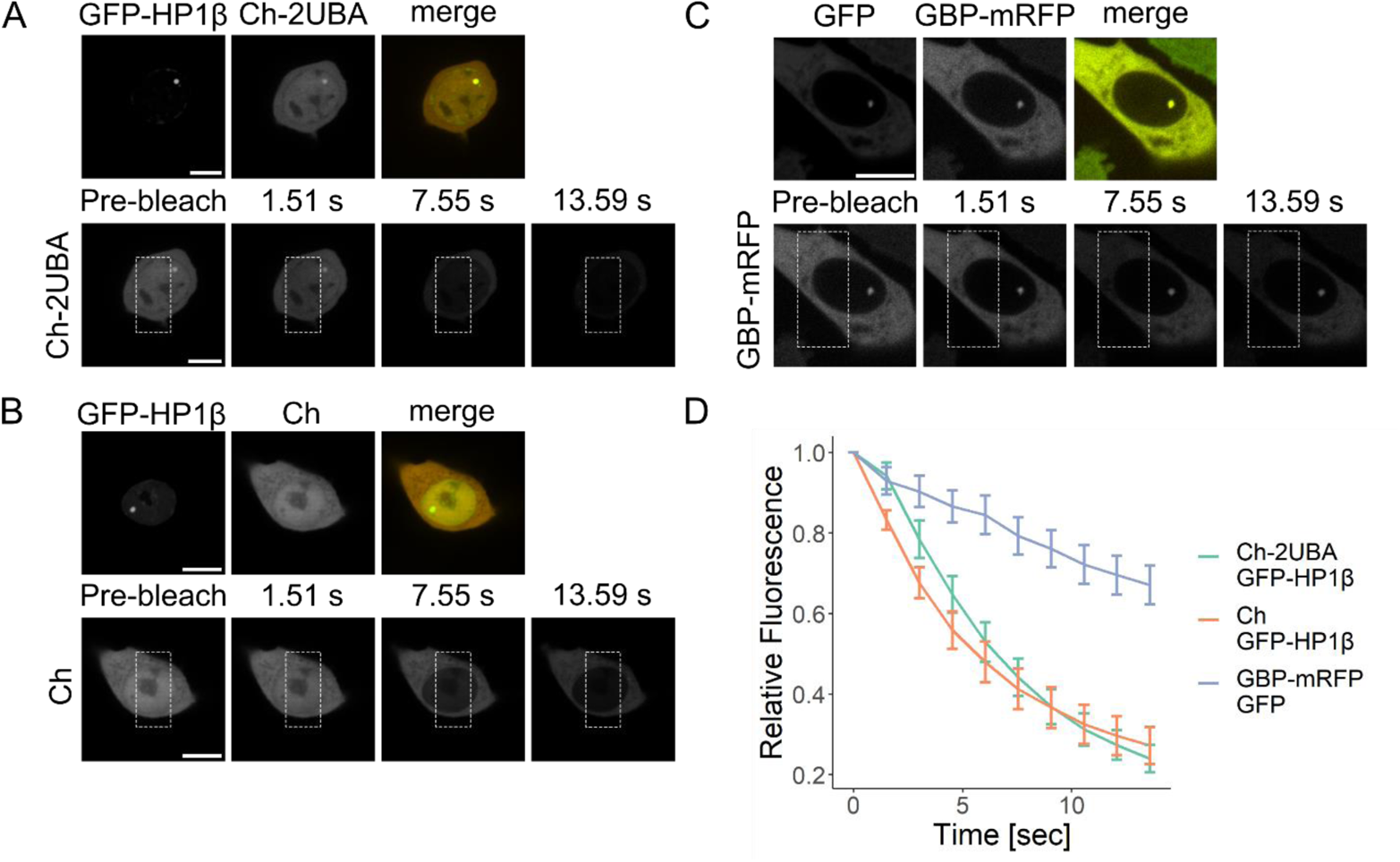
Related to Fig. 2B. Ch-2UBA transiently interacts with GFP-HP1β. The error bars depict the standard error of the mean. Ch-2UBA/GFP-HP1β n=13 (A), Ch/GFP-HP1β n=11 (B), GFP/GBP-mRFP n=18 (C). Scale bars = 10 µm.

**Extended Data Fig. 4:**
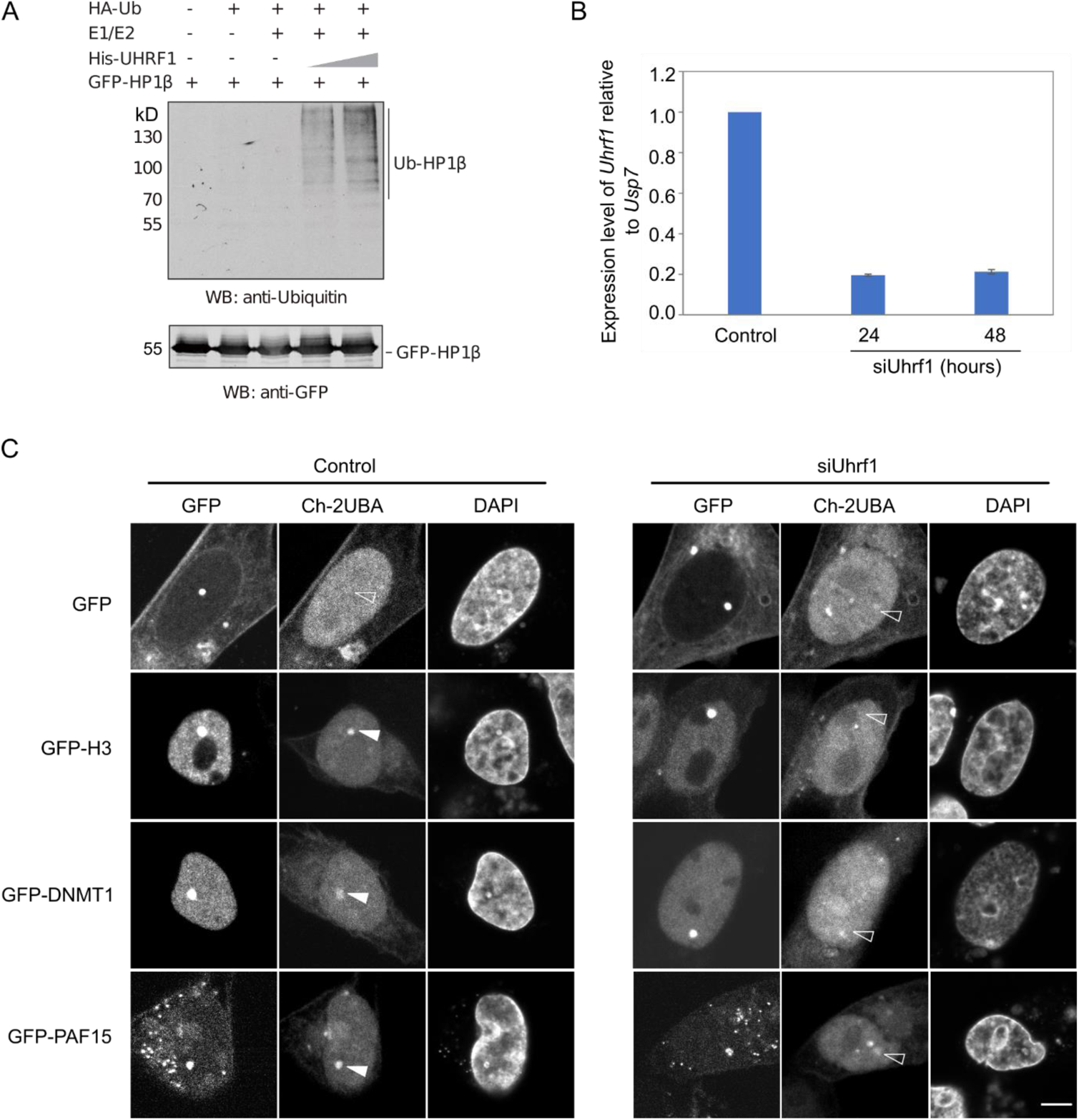
Related to Fig. 2B. Characterization of UHRF1 ubiquitin targets. (A) UHRF1 ubiquitinates HP1β *in vitro*. GFP-HP1β and His-UHRF1 were expressed and purified from HEK293T cells using the GFP-Trap and Ni-NTA beads, respectively. *In vitro* ubiquitination assays were performed with the components as indicated. Ubiquitination of HP1β was detected with an anti-Ubiquitin antibody. (B) *Uhrf1* mRNA levels in BHK wt and *Uhrf1*-knockdown cells were determined by quantitative RT-PCR. Expression data were normalized to *Usp7*. Relative mRNA levels are shown as mean values ± SEM from four independent technical repeats. (C) Representative images show the ubiquitination of DNMT1, H3 and PAF15 with the ubiF3Hc assay in BHK wt or *Uhrf1*-knockdown cells. The GFP fusion proteins were co-expressed with Ch-2UBA. Ch-2UBA accumulation at *lacO* spots reflects the ubiquitination of GFP fusion proteins. Scale bars: 5 µm.

**Extended Data Fig. 5:**
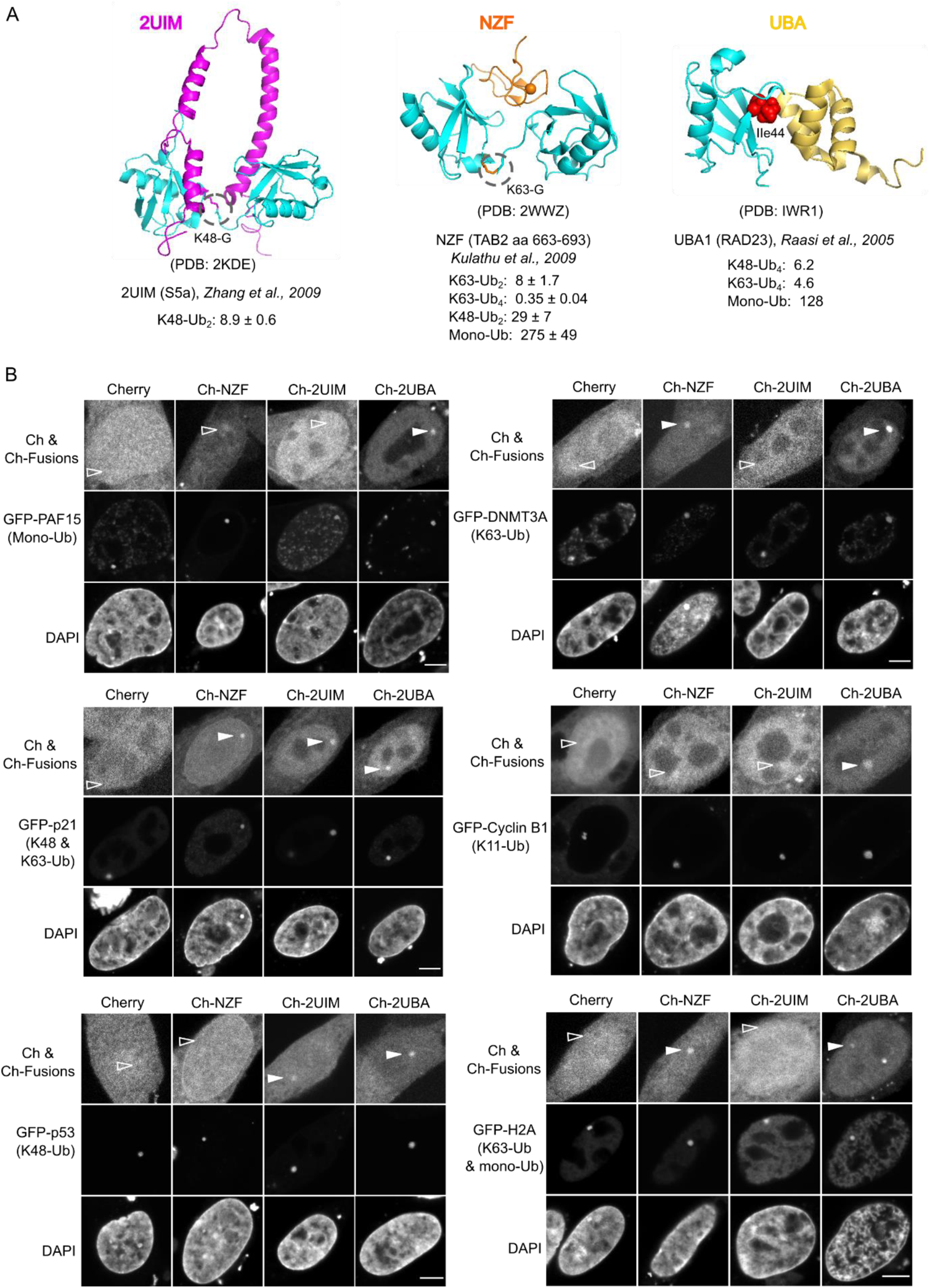
Related to Fig. 3B and 3C. (A) Structure of ubiquitin probes with ubiquitins and their reported *K*d values for ubiquitin interactions. 2UIM and K48-Ub2; NZF and K63-Ub2; UBA and Ub. 3D structures are from the Protein Data Base (PDB) as indicated. (B) The ubiquitination of GFP fusion proteins was detected by ubiF3H assays using different ubiquitin probes as indicated. Representative images are shown and ubiquitination of GFP fusion proteins detected with the different probes indicated is highlighted with filled arrowheads and not detectable ubiquitination is indicated with open arrowheads (for quantification see Extended Data Fig. 6). Scale bars: 5 µm.

**Extended Data Fig. 6:**
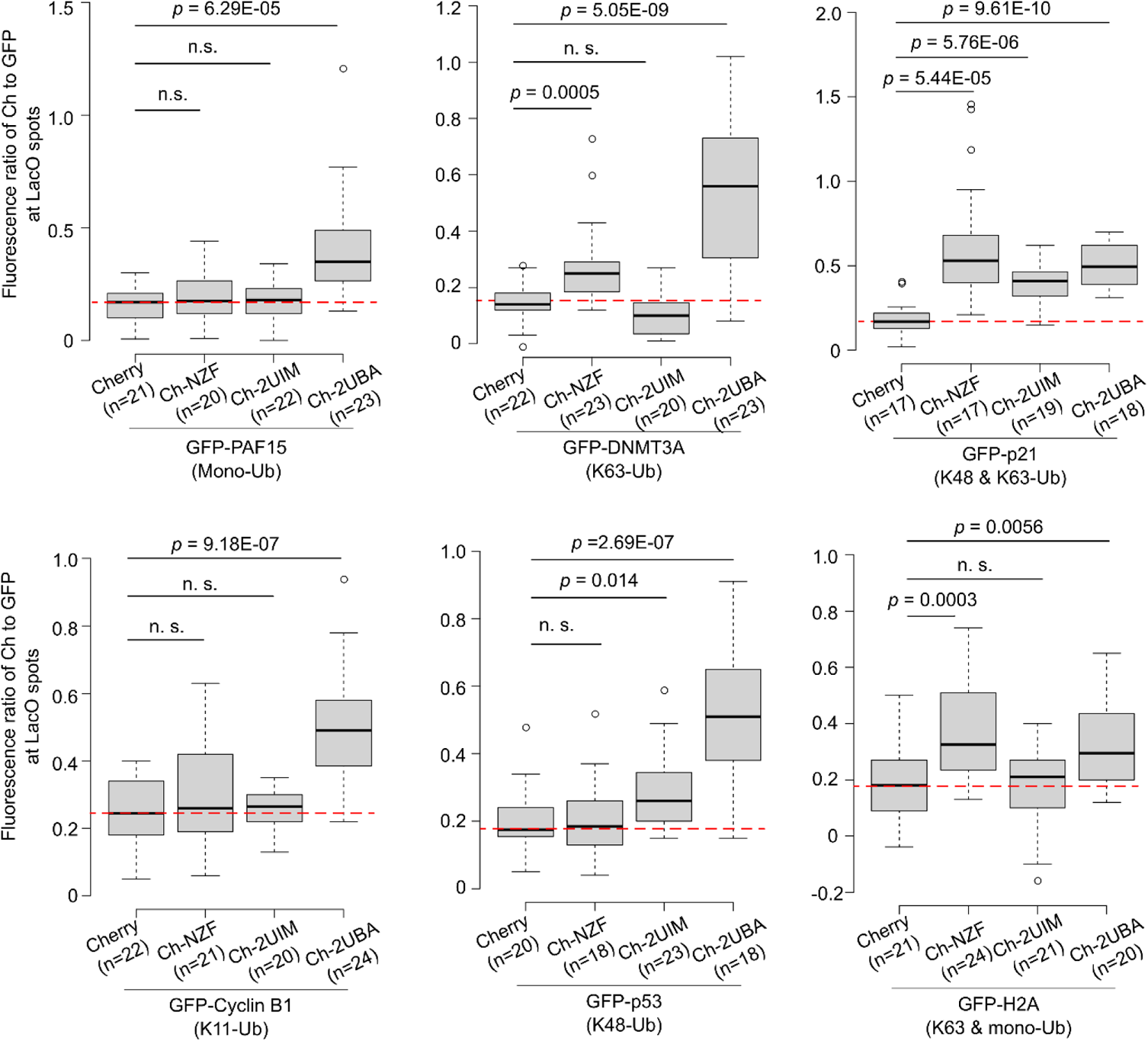
Related to Fig. 3C. The ubiquitination of GFP fusion proteins was detected by ubiF3H assays using different ubiquitin probes. Quantification of the intensity ratio of mCherry-tagged ubiquitin probes to GFP fusion proteins at the *lacO* array. Center lines show the medians; box limits indicate the 25th and 75th percentiles as determined by R software; whiskers extend 1.5 times the interquartile range from the 25th and 75th percentiles; outliers are represented by dots. Cells analyzed (n) are indicated. The red dot lines indicate the median of corresponding controls. Unpaired student t-tests were performed, and *p*-values indicated. The *p*-value greater than 0.05 is considered as not significant (n.s.).

**Extended Data Fig. 7:**
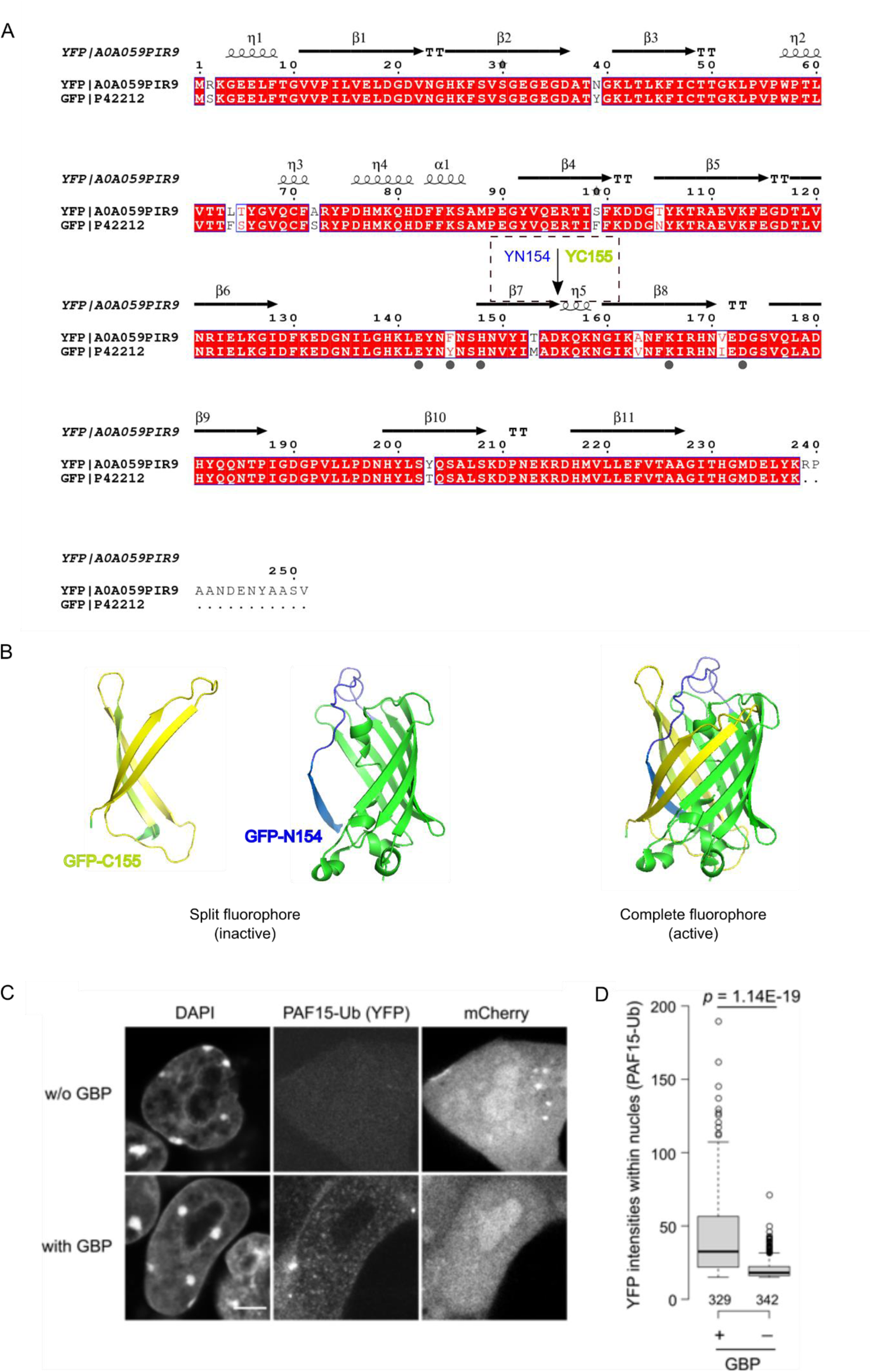
Related to Fig. 4. Split YFP for fluorescence complementation assay. (A) Protein sequence alignment of GFP and YFP using a website tool (http://espript.ibcp.fr/ESPript/ESPript/). The secondary structures are shown above. Secondary structure elements are: α, α-helices; η, 310-helices; β, β-strands; TT, strict β-turns. Strict sequence identity is shown by a red box with white characters. The split site (YN154/YC155) and the major amino acids of GFP or YFP in the interaction surface with GBP are highlighted with black dots beneath the sequences. (B) Structures show the inactive two halves of GFP, GFP- C155 indicated in yellow and GFP-N154 in green (left). The β-sheet 7 in GFP-N154 is labeled in blue. The active reconstituted GFP with GBP protein (PDB ID code 3K1K) is shown (right). (C) GBP enhances the reconstituted YFP signals. Representative images of detection of PAF15 ubiquitination by ubiF3Hc with or without GBP in mESCs (left). The expression of mCherry was used to identify transfected cells. Scale bar: 5 µm. (D) mESCs expressing Cherry, YC-PAF15 and 2UBA-YN with or without GBP were analyzed with the Harmony analysis software (PerkinElmer) and the intensities of reconstituted YFP signals in the nucleus were measured (right). Center lines show the medians; box limits indicate the 25th and 75th percentiles as determined by R software; whiskers extend 1.5 times the interquartile range from the 25th and 75th percentiles; outliers are represented by dots. Cells analyzed (n) and p value are indicated.

**Extended Data Fig. 8:**
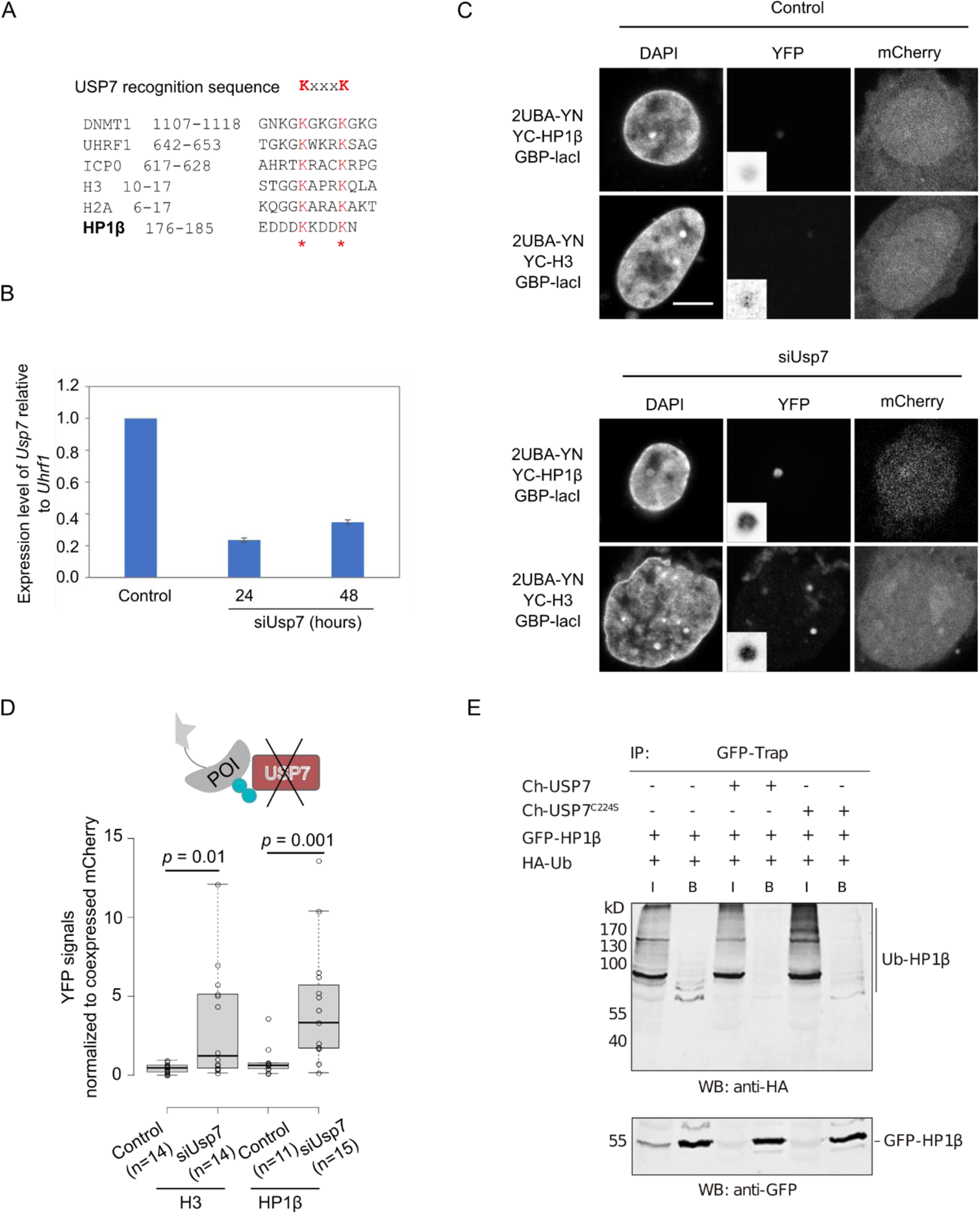
Related to Fig. 4. USP7 is the deubiquitinase for HP1β ubiquitination. (A) The alignment of the KxxxK motif of HP1β to the corresponding sequences of DNMT1, UHRF1, ICP0, H3 and H2A. Conserved lysine residues of the KxxxK motif are highlighted in red. (B) *Usp7* mRNA levels in BHK wt and *Usp7*-knockdown cells were determined by quantitative RT-PCR. Expression data are normalized to *Uhrf1*. Relative mRNA levels are shown as mean values ± SEM from four independent technical repeats. (C) Representative images of H3 (positive control) and HP1β ubiquitination with the ubiF3Hc assay in BHK wt and *Usp7*-knockdown cells in D. Scale bar: 5 µm. (D) Quantification of YFP intensities at *lacO* arrays normalized to co-expressed mCherry in BHK wt and *Usp7*-knockdown cells. Center lines show the medians; box limits indicate the 25th and 75th percentiles as determined by R software; whiskers extend 1.5 times the interquartile range from the 25th and 75th percentiles; outliers are represented by dots. The numbers of cells analyzed (n) are indicated. Unpaired student t-tests were performed, and *p*- values indicated. (E) Deubiquitination of HP1β by USP7. GFP-Trap pulldowns from HEK293T cells expressing indicated combinations of HA-Ub, GFP-HP1β, Ch-USP7 and Ch-USP7^C224S^ were determined by WB analysis with an anti-HA antibody.

**Extended Data Fig. 9:**
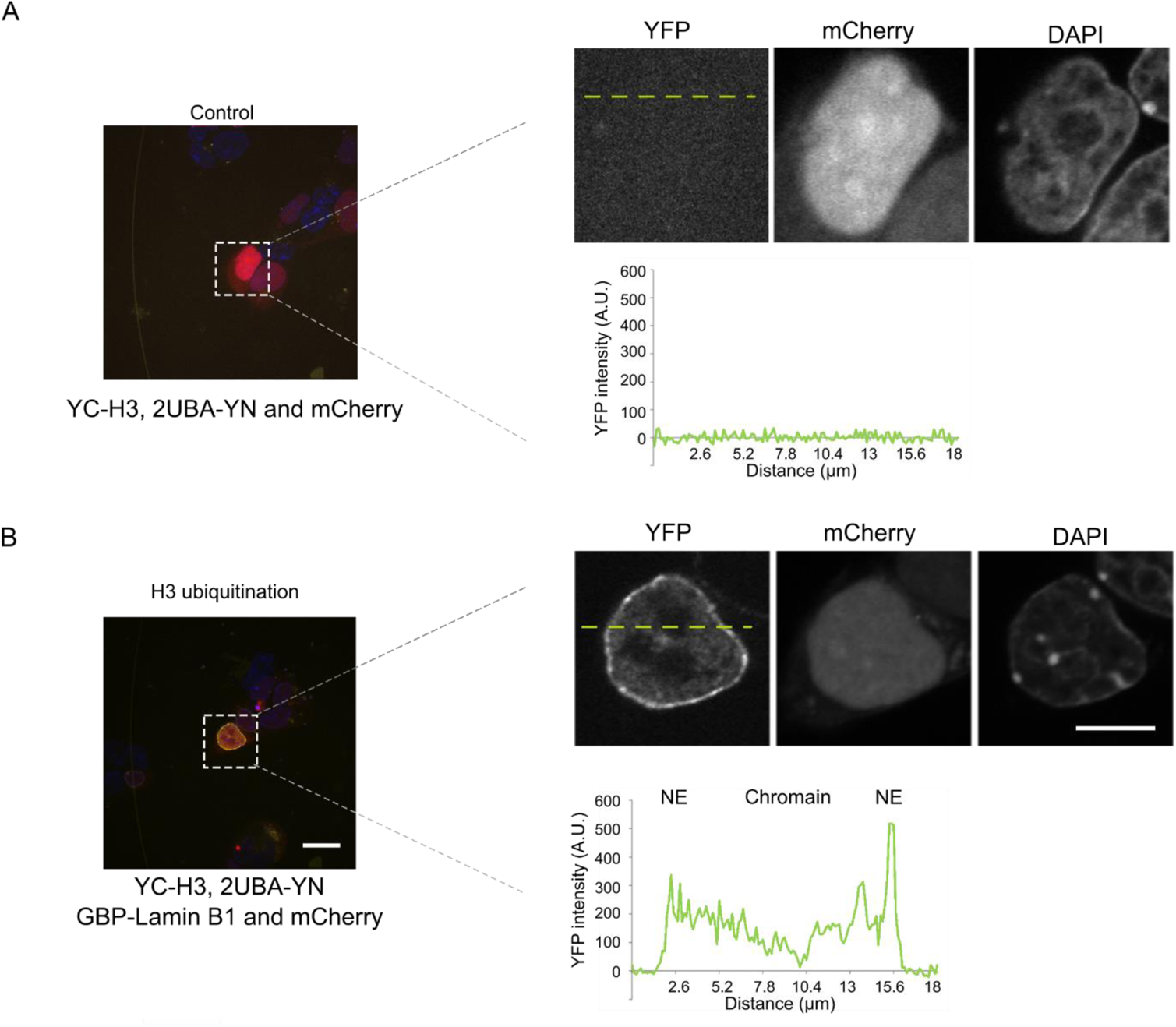
Related to Fig. 5B. Spinning disk confocal images of H3 ubiquitination with the ubiF3Hc assay. The ubiquitinated H3 complexes are trapped at the nuclear envelope by GBP-Lamin B1. Representative images of mESCs co-transfected with constructs as indicated in A (control without GBP) and B. The line plots show the intensity distribution of YFP along the scan line indicated in green. Scale bars: 20 µm for overview and 10 µm for zoomed in single cells.

**Extended Data Fig. 10:**
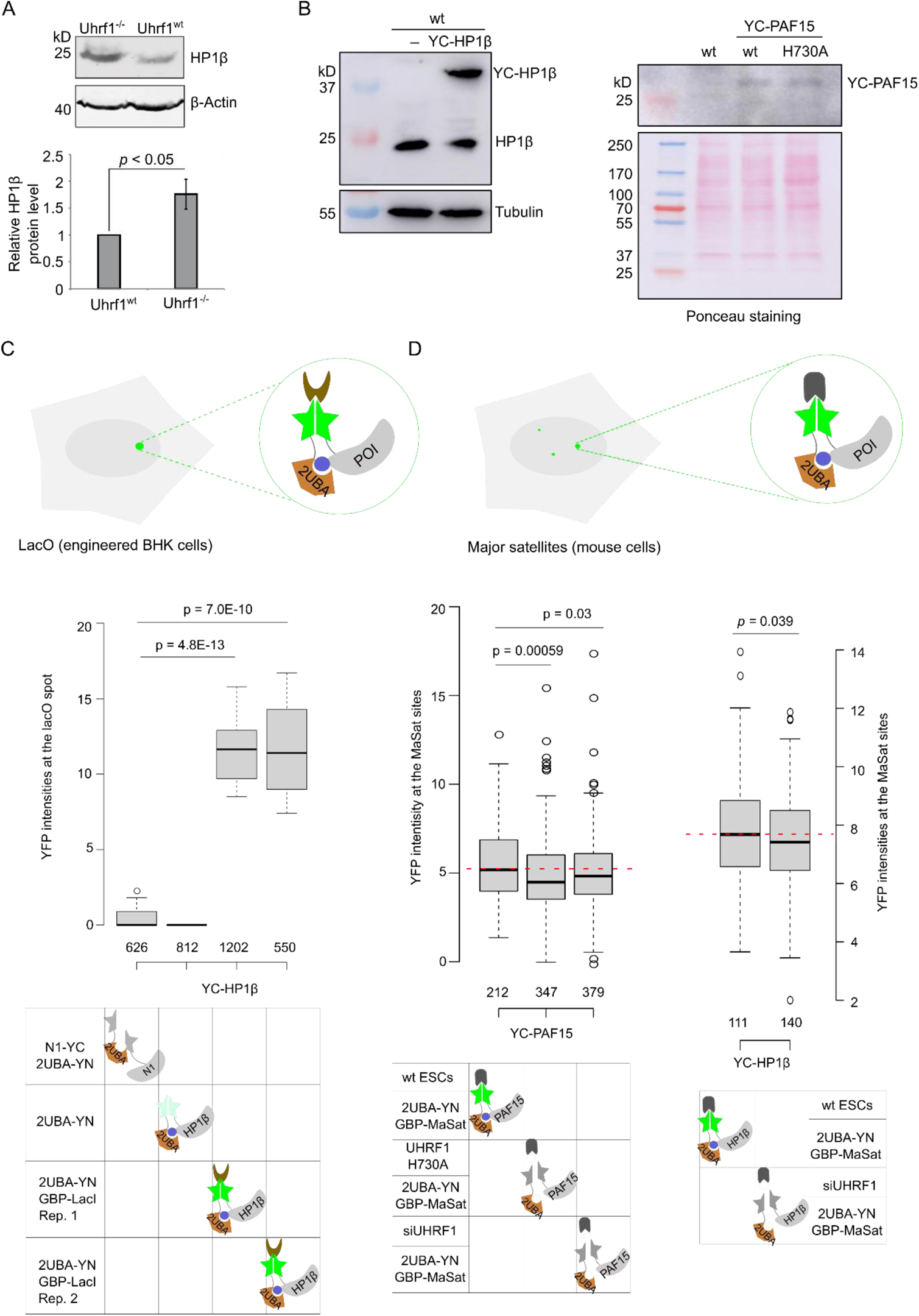
Related to Fig. 5F, 6B and 7. (A) HP1β levels are controlled by UHRF1. WB analysis of endogenous HP1β levels in wt and *Uhrf1*-deficient mESCs. An anti-β-actin blot was shown as loading control. Quantification of the HP1β protein levels was done with a gel analysis tool in ImageJ. Shown are mean values ± SEM of four independent biological replicates and normalized to HP1β level in wt mESCs. Differences in relative HP1β protein levels between wt and *Uhrf1*-deficient mESCs were analyzed using student’s t-test and considered statistically significant for *p* < 0.05 (*). (B) Characterization of mESC lines stably expressing YC-HP1β (left) or YC-PAF15 (right) by western blot. The anti-tubulin blot and Ponceau staining were used as loading controls. (C and D) Detection of protein ubiquitination with ubiF3Hc in 96-well plates. Center lines show the medians; box limits indicate the 25th and 75th percentiles as determined by R software; whiskers extend 1.5 times the interquartile range from the 25th and 75th percentiles; outliers are represented by dots. The numbers of cells analyzed are indicated. Data sets were tested for significance with an unpaired t-test and *p*-values are indicated. Detection of HP1β ubiquitination with ubiF3Hc by recruiting to lacO spots. Box plot representations of YFP intensities at *lacO* spots normalized to mCherry signals in the nucleus (C). Detection of HP1β and PAF15 ubiquitination in mESCs by recruiting to major satellite sites (MaSat). The red dot lines indicate the median of corresponding controls (D).

